# Nanochiral graphene quantum dots preferentially target virus-organized membrane states for host-sparing antiviral activity

**DOI:** 10.64898/2026.09.26.754715

**Authors:** Yichen Liu, Mahnoosh Maleki, Pilar Pérez-Romero, Yichun Wang

## Abstract

Selectively perturbing pathogenic membranes while preserving host-cell integrity is a fundamental challenge in the design of membrane-active biomaterials. This challenge is particularly acute for enveloped viruses because their lipid membranes are derived from host cells and therefore share many of the same molecular constituents. Here, we demonstrate that engineered nanochirality provides a structural parameter for distinguishing virus-organized membrane nanodomains from host-cell membranes. Histidine-functionalized graphene quantum dots (His-GQDs) form enantiomeric nanoscale structures with distinct interfacial topologies despite closely matched size, composition, and surface charge. *D*-His-GQDs preferentially interact with cholesterol-rich ordered membrane nanodomains associated with S-acylated coronavirus Spike, penetrate more deeply into raft-like lipid bilayers, and induce lipid disordering, membrane leakage, and viral-envelope disruption. Reducing Spike *S*-acylation or depleting membrane cholesterol attenuated this stereoselective interaction, while molecular simulations revealed asymmetric lipid organization surrounding *S*-acylated Spike and preferential insertion of *D*-His-GQDs into the ordered membrane environment. Functionally, *D*-His-GQDs directly inactivated human coronavirus OC43 (HCoV-OC43), inhibited infection with an EC50 of 0.92 μg mL□^1^ and exhibited a selectivity index of 361. Stereoselective antiviral activity extended to human cytomegalovirus, demonstrating activity across distinct enveloped-virus families, and intranasal *D*-His-GQDs protected HCoV-OC43-challenged mice under both prophylactic and early post-exposure regimens. Overall, these findings establish structural nanochirality as a biomaterial design parameter for recognizing higher-order membrane organization and converting virus-associated membrane states into selectively addressable therapeutic vulnerabilities.

## 1. Introduction

Enveloped viruses remain major drivers of recurrent epidemics and zoonotic spillover, highlighting the need for rapidly deployable antivirals when vaccines are unavailable or provide incomplete protection [1–4]. Most approved antivirals target specific viral proteins to block attachment, entry, or membrane fusion, or inhibit intracellular enzymes involved in genome replication, protein processing, and virion maturation [5–7]. Although highly potent, these antivirals target mutable viral components, enabling resistance to erode efficacy and forcing each emerging virus into a new cycle of target identification, drug development, and resistance surveillance rather than leveraging a conserved viral feature [8]. The lipid envelope represents one such conserved feature shared across many clinically important viruses. Acquired from host membranes, the viral envelope is essential for virion stability, cellular entry, and membrane fusion across diverse viral families [9, 10]. Targeting this shared feature could therefore enable cross-family antiviral activity while imposing a higher barrier to classical sequence-based resistance [11, 12]. However, the same host-derived origin that makes the viral envelope broadly conserved also poses a fundamental challenge for membrane-active material design: achieving selective targeting of viral membranes while sparing host cell membranes [13, 14]. Accordingly, conventional membrane-active materials relying on nonspecific amphipathic insertion or lipid perturbation often show increasing host-membrane toxicity with greater antiviral activity [15–18]. Overcoming this trade-off requires materials capable of recognizing differences in membrane organization rather than simply differences in membrane composition.

This selectivity challenge reflects an incomplete view of viral envelopes as miniature host membranes distinguished only by bulk lipid composition, net charge, or amphiphile susceptibility [19, 20]. Viral envelopes, however, are not simply small fragments of host membranes. Viral assembly and budding reorganize host-derived lipids and proteins into highly curved, densely crowded, and frequently cholesterol- and sphingolipid-enriched domains [21–23]. Repetitive viral glycoproteins, covalent lipid modifications, leaflet asymmetry, and restricted lateral remodeling can create an interfacial physical state that differs from the corresponding producer-cell membrane [24, 25]. This distinction is amplified after viral release: extracellular virions cannot actively traffic, remodel, or repair their envelopes, allowing localized membrane defects to propagate into irreversible loss of infectivity [9, 26]. Thus, viral and host membranes may share molecular constituents while exhibiting substantially different nanoscale interfacial states and capacities to recover from perturbation. These higher-order differences, including membrane order, lateral phase behavior, and protein–lipid nanodomain organization, are increasingly recognized as selective biophysical cues [20, 27–29]. A useful precedent comes from bacterial membranes, where accessible anionic surfaces and distinct lipid-packing environments enable amphipathic cationic peptides to preferentially bind and induce pore formation or non-lytic membrane destabilization [30, 31]. Hence, we reasoned that the distinct physical states of viral and host membranes could provide a basis for selective recognition by appropriately structured nanomaterials.

Structural nanochirality provides a potential means to interrogate such membrane organization. At nano–bio interfaces, enantiomeric materials can exhibit significantly different adhesion, penetration, and membrane-perturbation behaviors even when their composition, size, and charge are otherwise matched [32–35]. Because matched nano-enantiomers present distinct 3D interfacial topologies without requiring major changes in bulk physicochemical properties, structural nanochirality may therefore function as a “selectivity dial” for nanoscale lateral organization, lipid-packing defects, and local membrane geometry [36–38]. Coronavirus Spike provides a mechanistically defined example of such a virus-organized membrane state: Spike *S*-acylation is critical for virion assembly and infectivity [39, 40] and promotes ordered, cholesterol- and sphingolipid-enriched nanodomains at viral assembly and budding sites [23, 41]. Chirality-dependent lipid packing, interfacial hydration, membrane curvature, and leaflet asymmetry may further impart local structural asymmetry [42–45], potentially creating an interface that can be differentially recognized by nano-enantiomers. Graphene quantum dots (GQDs) provide a versatile platform for testing this principle because of their high surface area, favorable biocompatibility, and readily programmable edge chemistry [46–48]. We previously showed that chiral ligand functionalization can induce out-of-plane distortion of GQDs and that these structurally chiral nanomaterials exhibit enantioselective transport across biological lipid membranes [49, 50]. These observations suggested that ligand-encoded GQD topology could be used not only to tune membrane affinity, but also to discriminate between distinct membrane physical states.

Here, we test this principle using *L*- and *D*-histidine-functionalized graphene quantum dots (*L*- and *D*-His-GQDs) as matched nano-enantiomeric biomaterials. Histidine edge functionalization generates handed distortions of the GQD framework and corresponding differences in interfacial topology [49, 50]. We hypothesized that these structural differences would produce stereoselective engagement of virus-organized ordered membrane nanodomains and thereby separate antiviral membrane disruption from host-cell toxicity. Using HCoV-OC43 as a mechanistically tractable enveloped-virus model, we show that *D*-His-GQDs preferentially associate with *S*-acylation- and cholesterol-dependent ordered membrane environments, penetrate more deeply into model bilayers, disrupt lipid order, and compromise envelope integrity. This stereoselective interaction directly reduces extracellular virion infectivity and infection-associated cell–cell fusion. *D*-His-GQDs inhibit HCoV-OC43 with an EC50 of 0.92 μg mL□^1^ and a selectivity index of approximately 361, while stereoselective activity extends to human cytomegalovirus in two human cell types. Intranasal administration further protects HCoV-OC43-challenged mice under prophylactic and early post-exposure treatment. Overall, these results establish a new design principle whereby engineered nanochirality selectively recognizes and disrupts virus-organized membrane states, providing a general biomaterials strategy for enhancing the therapeutic selectivity of membrane-active nanomaterials.

**Fig. 1.**
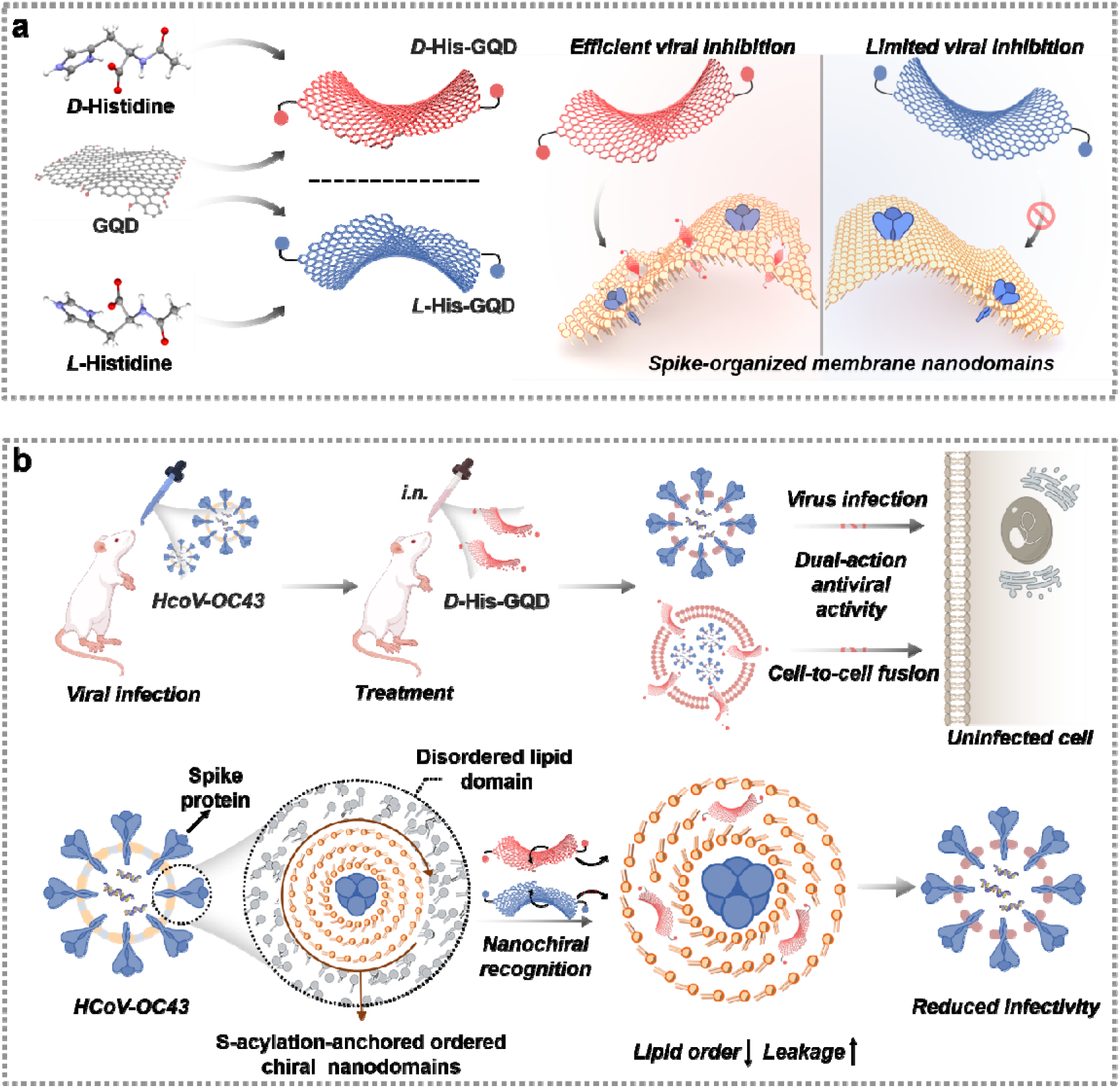
Design and proposed dual antiviral action of histidine-functionalized graphene quantum dots. (a) Synthesis of enantiomeric *L*- and *D*-histidine-functionalized graphene quantum dots (*L*- and *D*-His-GQDs) and their differential interactions with Spike-associated ordered membrane nanodomains. (b) Proposed model in which *D*-His-GQDs directly inactivate extracellular human coronavirus OC43 (HCoV-OC43) virions and attenuate infection-associated syncytium formation through preferential interaction with ordered membrane nanodomains organized around S-acylated Spike.

## 2. Results and discussion

### 2.1. Synthesis and characterization of chiral GQDs

*L*- and *D*-His-GQDs were synthesized by modifying pristine GQDs with *L*- or *D*-histidine through 1-ethyl-3-(3-dimethylaminopropyl)carbodiimide hydrochloride/N-hydroxysuccinimide (EDC·HCl/NHS)-mediated coupling, following our established protocol (Fig. 2a) [49]. Fourier transform infrared spectroscopy supported successful conjugation through the appearance of a C–N stretching band at approximately 1250 cm^-1^ (Fig. 2b). ζ-potential analysis further revealed a shift from −21.3 ± 1.2 mV for carboxyl-rich pristine GQDs to +1.7 ± 0.4 and +2.4 ± 0.3 mV for *L*- and *D*-His-GQDs, respectively (Fig. 2c). This is consistent with consumption of edge carboxyl groups during amide formation and a change toward a slightly cationic surface. UV–vis absorption spectra of *L*- and *D*-His-GQDs showed a red shift of the π–π* band to 262 nm relative to pristine GQDs (Fig. S1). Consistently, the photoluminescence spectra displayed the maximum emission shifting from ∼420 nm to ∼450 nm (Fig. 2d). These spectral shifts suggest electronic coupling between the aromatic GQD framework and histidine-derived surface states, with possible contributions from ***n***–π* transitions. Circular dichroism (CD) spectroscopy showed that pristine GQDs were chiroptically silent, whereas *L*- and *D*-His-GQDs exhibited mirror-image Cotton effects centered at 258 nm, coincident with their UV–vis absorption bands (Fig. 2e). These signals support transfer of ligand chirality to the chiroptical response of the functionalized GQDs. Transmission electron microscopy (TEM) showed that *L*- and *D*-His-GQDs retained lateral dimensions of approximately 10–13 nm, comparable to those of pristine GQDs (Fig. 2f). High-resolution TEM revealed lattice fringes with an interplanar spacing of 0.23 nm, assigned to the graphene (112 0) plane. However, unlike the continuous lattice of pristine GQDs, the chiral GQDs exhibited multiple localized distortions. Atomic force microscopy further characterized this change in surface topography (Fig. 2g). Pristine GQDs showed a topographic height of ∼0.4 nm, consistent with monolayer graphene. In contrast, *L*- and *D*-His-GQDs exhibited an increased thickness ranging from 0.6 to 0.9 nm. The increased apparent height is consistent with histidine functionalization at the GQD edges and the resulting change in surface topography. To examine the atomistic basis of the observed chiroptical response, density functional theory (DFT) geometry optimizations were performed at the B3LYP/6-31G(d) level. Geometry optimization revealed that histidine functionalization induces a saddle-shaped out-of-plane deformation of the GQD framework, with *L*- and *D*-His-GQDs adopting mirror-related handed topologies (Fig. S2). Functionalization-induced redistribution of interaction regions was visualized using interaction region indicator maps. In contrast to pristine GQDs, which exhibit a highly symmetric and spatially uniform hexagonal pattern, both *D*- and *L*-His-GQDs displayed pronounced symmetry breaking, anisotropic redistribution of interaction regions, and disrupted continuity of the π-conjugated network, particularly near the functionalized edges (Fig. 2h). Electrostatic potential analysis yielded a molecular polarity index (MPI) of 14.31 kcal mol□^1^ for pristine GQDs (Fig. 2i and Fig. S3). However, MPI values for the *L*- and *D*-His-GQD were 5.57 and 4.72 kcal mol^−1^, respectively, suggesting a shift toward edge-localized interaction sites. Reduced density gradient analysis revealed extensive hydrogen-bonding interactions between the imidazole and amino groups of histidine and oxygen-containing groups at the GQD edges, which may contribute to the observed distortion (Fig. S4). Time-dependent DFT calculations reproduced the qualitative mirror-image Cotton effects observed experimentally (Fig. S5). These experimental and computational results support handedness-dependent differences in the interfacial organization of *L*- and *D*-His-GQDs (Fig. 2j), providing a structural basis for testing their stereoselective interactions with biological membranes.

**Fig. 2.**
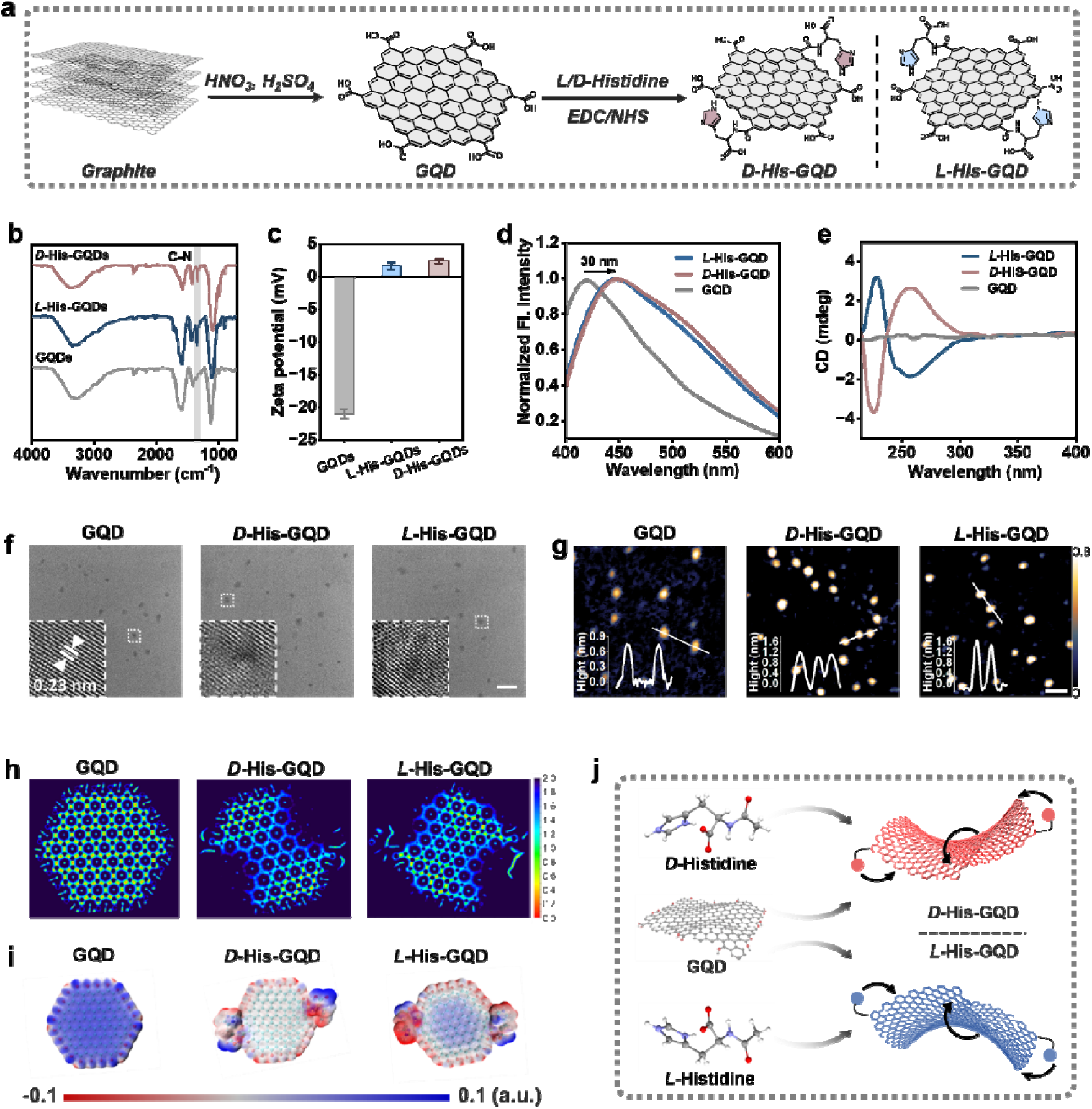
Synthesis and structural characterization of histidine-functionalized graphene quantum dots (His-GQDs). (a) Synthesis of pristine graphene quantum dots (GQDs) and conjugation with *L*- or *D*-histidine through 1-ethyl-3-(3-dimethylaminopropyl)carbodiimide/N-hydroxysuccinimide (EDC/NHS) coupling. (b) Fourier-transform infrared spectra of pristine GQDs, *L*-His-GQDs, and *D*-His-GQDs. (c) ζ-potentials of the indicated GQDs. Data are mean ± SD (*n* = 3). Normalized photoluminescence emission spectra (d) and circular dichroism (CD) spectra (e) of pristine GQDs and *L*- and *D*-His-GQDs. (f) Transmission electron microscopy (TEM) images with high-resolution TEM (HRTEM) images shown in the insets. Scale bars, 20 nm. (g) Atomic force microscopy images and corresponding height profiles. Scale bars, 50 nm. Interaction region indicator maps (h) and electrostatic potential maps (i) calculated for the density functional theory (DFT)-optimized structures of pristine GQDs and *L*- and *D*-His-GQDs. (j) Proposed saddle-shaped conformations of *L*- and *D*-His-GQDs and their mirror-related nanochiral interfaces.

### 2.2. Nanochirality confers enantioselective activity against enveloped viruses

Antiviral activity was evaluated against HCoV-OC43 using remdesivir and CPC as mechanistically distinct controls. Remdesivir inhibits intracellular RNA polymerase activity, whereas CPC is a cationic membrane-active virucide that disrupts lipid-enveloped viruses (Fig. S6) [51–54]. In plaque assays, *D*-His-GQDs nearly abolished plaque formation, while pristine GQDs and *L*-His-GQDs had little effect. CPC and remdesivir also reduced plaque numbers, but residual plaques remained more evident than with *D*-His-GQDs (Fig. S7). *D*-His-GQDs suppressed infectious HCoV-OC43 production in a concentration-dependent manner, with an EC50 of 0.92 μg mL^-1^, compared with 1.08 μg mL^-1^ for remdesivir and 1.86 μg mL^-1^ for CPC (Fig. 3a). The distinction was more pronounced at the cellular level: *D*-His-GQDs exhibited CC50 values of 207.9 ± 8.0, 146.6 ± 3.4, and 332.0 ± 4.4 μg mL^-1^ in SH-SY5Y, HepG2, and RD cells, exceeding the corresponding CPC values by approximately 189-, 73-, and 98-fold (Fig. S8). These results show that *D*-His-GQDs combine antiviral potency with a substantially wider host-cell tolerance window than the conventional membrane-active controls. In RD cells, the resulting selectivity index was approximately 361 for *D*-His-GQDs, compared with 14.3 for remdesivir and 1.8 for CPC. During multicycle infection, *D*-His-GQDs maintained infectious titers near 3 log_10_ TCID_50_ mL^-1^ through 72 h, whereas titers reached approximately 3.6 log_10_ with remdesivir and 4.5 log_10_ with CPC. Untreated cultures and those receiving pristine or *L*-His-GQDs approached 6 to 7 log_10_ TCID_50_ mL^-1^ (Fig. S9). Thus, *D*-His-GQDs produced the most sustained suppression under continuous treatment. A time-of-contact assay distinguished direct virion inactivation from intracellular replication inhibition. *D*-His-GQDs produced a progressive loss of infectivity that reached approximately 2.8 log_10_ after 90 min, compared with approximately 2.2 log_10_ for CPC, whereas remdesivir, pristine GQDs, and *L*-His-GQDs showed little direct activity (Fig. 3b). The stage-of-action profile further separated the membrane-active compounds from remdesivir. *D*-His-GQDs and CPC were effective during full-time and entry-only exposure but showed little post-entry activity, whereas remdesivir acted predominantly after entry (Fig. S10). Consistently, virions preincubated with the indicated agents at 10 μg mL^-1^ for 1 h at 37 °C produced distinct cellular Spike signals, with *D*-His-GQDs yielding the lowest signal, followed by CPC, whereas the other GQD formulations remained largely ineffective (Fig. 3c and Fig. S11). At 10 μg mL^-1^, residual infectivity after *D*-His-GQD treatment was approximately 6.1% at 37 °C and 64.5% at 4 °C, suggesting that temperature-dependent membrane dynamics contribute to virucidal activity (Fig. S12). Because HCoV-OC43 Spike can promote cell-cell fusion and facilitate direct spread between neighboring cells [55, 56], we examined syncytium formation under authentic infection conditions.

**Fig. 3.**
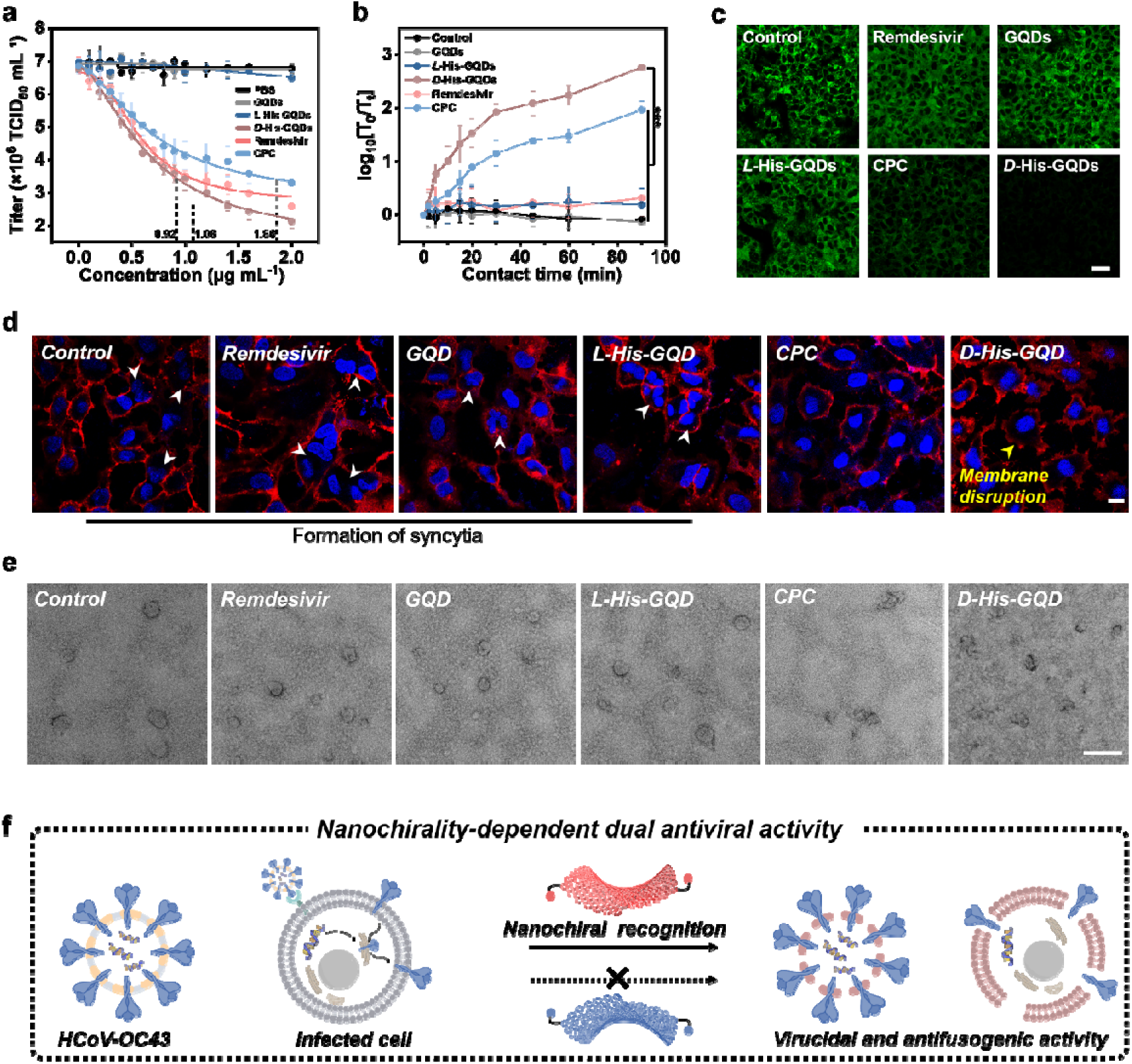
Enantioselective inhibition and direct inactivation of human coronavirus OC43 (HCoV-OC43) by *D*-His-GQDs. (a) Concentration-dependent effects of pristine GQDs, *L*-His-GQDs, *D*-His-GQDs, remdesivir, and cetylpyridinium chloride (CPC) on infectious HCoV-OC43 production in human rhabdomyosarcoma (RD) cells at 48 h. Dashed lines indicate half-maximal effective concentration (EC) values of 0.92, 1.08, and 1.86 μg mL□^1^ for *D*-His-GQDs, remdesivir, and CPC, respectively (*n* = 5). (b) Contact-time-dependent virucidal activity after incubation of HCoV-OC43 virions with the indicated agents at 10 μg mL^-1^ for 0 to 90 min. T_0_ and T_t_ denote infectious titers at 0 and t min, respectively, and log_10_(T_0_/T_t_) represents the reduction in infectious titer (*n* = 5). (c) Representative immunofluorescence images of RD cells challenged with compound-treated virions. Virions were preincubated with the indicated compounds at 10 μg mL^-1^ for 1 h at 37°C before challenge. Scale bar, 25 μm. (d) Representative fluorescence images of infected RD cells under the indicated treatments. Cell membranes are in red and nuclei in blue. Arrowheads indicate multinucleated syncytia. Scale bar, 10 μm. (e) TEM images of virions after incubation with the indicated materials. Scale bar, 200 nm. (f) Proposed model of nanochirality-dependent extracellular virion inactivation and reduction of infection-associated syncytia. Data are mean ± SD. Statistical comparisons were performed using two-tailed *t* tests. \*\*\**P* < 0.001.

Extensive syncytia were present in untreated cultures and remained evident after treatment with pristine GQDs or *L*-His-GQDs. *D*-His-GQDs markedly reduced syncytial morphology and produced altered membrane profiles in infected RD cultures (Fig. 3d). This response was accompanied by increased lactate dehydrogenase (LDH) release (Fig. S13), despite the high tolerance of uninfected RD cells to *D*-His-GQDs (Fig. S8). CPC also reduced syncytial morphology, but its much lower RD-cell CC_50_ indicates substantial overlap between its antiviral and host-cell membrane activities. TEM further revealed disrupted virion morphology after exposure to either *D*-His-GQDs or CPC, whereas virions treated with pristine GQDs, *L*-His-GQDs, or remdesivir largely retained their spherical appearance (Fig. 3e). These findings show that *D*-His-GQDs and CPC can physically compromise extracellular virions, while *D*-His-GQDs couple this membrane activity to a broad cellular tolerance window (Fig. 3f). The potential extension of chirality-dependent antiviral activity across enveloped virus families was evaluated using HCMV, a phylogenetically distant enveloped DNA virus with broad cellular tropism. We evaluated *L*- and *D*-His-GQDs in MRC-5 human lung fibroblasts and ARPE-19 human retinal pigment epithelial cells infected with BADrUL131-Y4 [57], a GFP-expressing derivative of HCMV strain AD169 in which the UL131 locus has been repaired. In time-of-addition assays, virus preincubation and cotreatment with *D*-His-GQDs markedly reduced infection in both MRC-5 and ARPE-19 cells at 48 and 72 h post-infection. Post-inoculation treatment remained partially effective, whereas *L*-His-GQDs showed little activity across all treatment schedules (Fig. S14 and S15). *D*-His-GQDs also inhibited HCMV infection in a concentration-dependent manner, with half-maximal inhibitory concentration (IC_50_) values of 8.39 μg mL^-1^ in MRC-5 cells and 1.04 μg mL^-1^ in ARPE-19 cells (Fig. S16). Both enantiomers exhibited favorable and comparable cytocompatibility, maintaining approximately 80% or greater viability after 24 h exposure at concentrations up to 10 μg mL^-1^ (Fig. S17), further supporting the stereoselective nature of the antiviral response. Thus, the stereoselective inhibition of phylogenetically distinct enveloped viruses supports the potential of *D*-His-GQDs as a broad-spectrum antiviral platform targeting early membrane-dependent stages of infection.

### 2.3. *D*-His-GQDs preferentially interact with membrane domains organized by Spike S-acylation

To test whether ordered membrane nanodomains of enveloped viruses constitute stereochemically addressable targets for nanochirality-dependent membrane disruption, we established a cell-surface Spike display model by plasmid transfection to assess the preferential association of *D*-His-GQDs with Spike-associated ordered (raft-like) membrane nanodomains, circumventing the technical challenges of directly resolving these nanoscale domains on individual virions. RD cells were transiently transfected to express wild-type (WT) HCoV-OC43 Spike or an eight-Cys-to-Ala mutant (Mut) with reduced Spike S-acylation (Fig. 4a). Non-permeabilized immunofluorescence imaging showed comparable cell-surface expression for both Spike variants (Fig. 4b and S18). Next, we imaged membrane order using a Nile Red-based solvatochromic (NR12S) probe, whose emission is blue-shifted in raft-like liquid-ordered (Lo) membranes (Fig. S19).

**Fig. 4.**
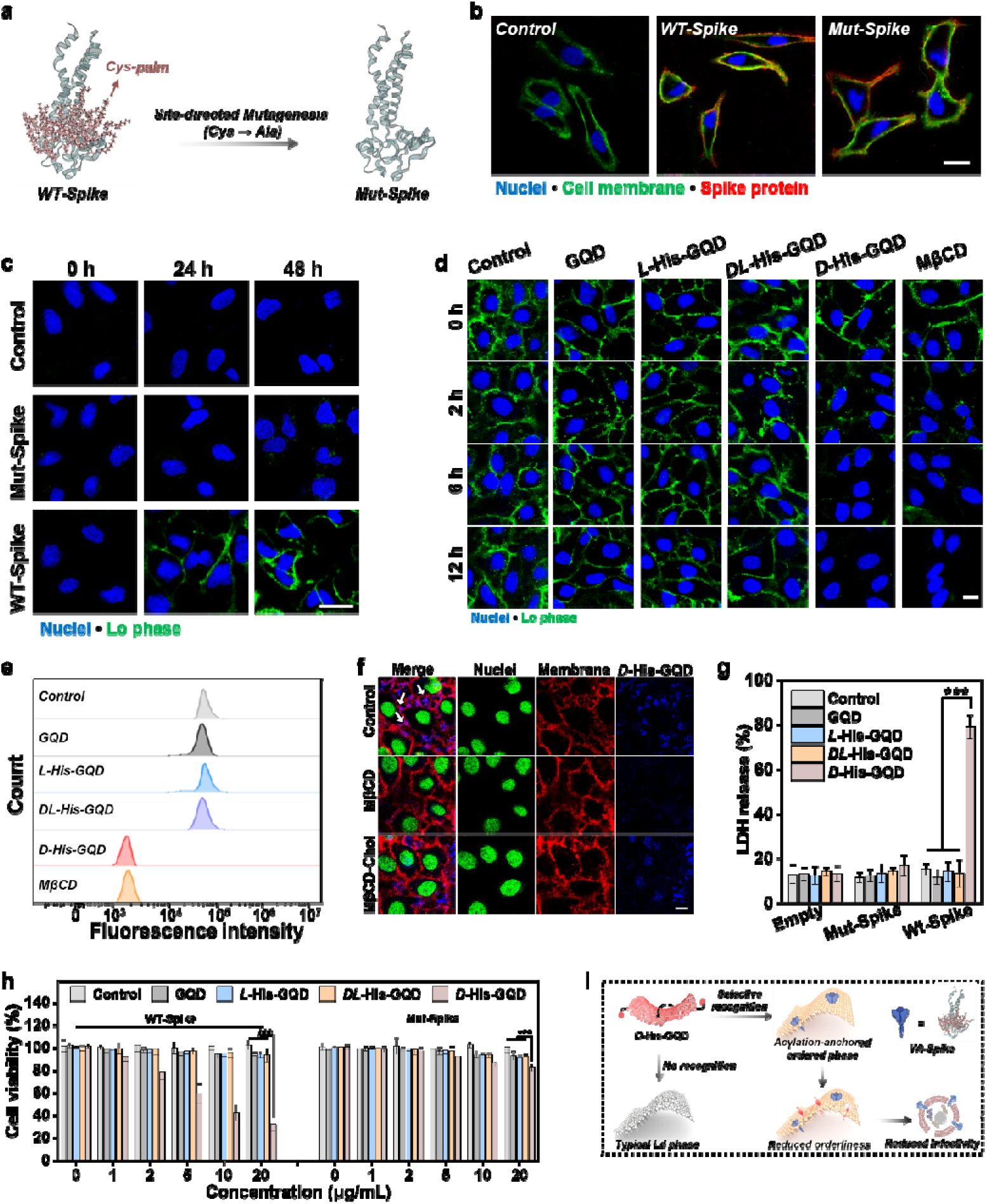
*D*-His-GQDs selectively perturb Spike-associated ordered membrane domains. (a) Schematic of human rhabdomyosarcoma (RD) cells expressing wild-type (WT) HCoV-OC43 Spike or an eight-Cys-to-Ala mutant (Mut) designed to reduce Spike S-acylation. (b) Representative non-permeabilized immunofluorescence images of surface WT and Mut Spike. Scale bar: 20 μm. (c) NR12S images of control, Mut-Spike, and WT-Spike cells at indicated times after transfection. Scale bar: 20 μm. (d) Time-dependent NR12S imaging of WT-Spike cells treated with the indicated GQDs or methyl-β-cyclodextrin (MβCD). Scale bar: 20 μm. (e) Flow-cytometric histograms of NR12S fluorescence after the indicated treatments. (f) *D*-His-GQD-associated fluorescence in WT-Spike cells following MβCD-mediated cholesterol depletion or repletion with cholesterol-loaded MβCD (MβCD–Chol). Scale bar: 10 μm. (g) Lactate dehydrogenase (LDH) release from empty-vector, Mut-Spike, and WT-Spike cells. (h) Viability of WT-Spike- and Mut-Spike-expressing cells after the indicated GQD treatments. (i) Proposed model of *D*-His-GQD recognition and disruption of Spike-associated ordered membrane domains. Data are mean ± SD (*n* = 5). Statistical comparisons were performed using two-tailed *t* tests. \*\*\**P* < 0.001.

WT-Spike-expressing cells displayed higher Lo-associated NR12S mean fluorescence intensity than control and Mut-Spike cells (Fig. S20), and the signal increased over time after WT-Spike transfection (Fig. 4c). The GQD variants produced distinct effects on the Lo-associated NR12S signal in WT-Spike-expressing cells. *D-*His-GQDs significantly reduced this signal within 2 h and brought it to near-background levels by 12 h, with an endpoint effect comparable to that of methyl-β-cyclodextrin (MβCD), a cholesterol-depleting agent commonly used to perturb ordered membrane domains (Fig. 4d and S21). Cholesterol repletion with cholesterol-loaded MβCD (MβCD–Chol) restored the Lo-associated NR12S signal in a concentration-dependent manner, with quantification confirming recovery at 100 μM (Fig. S22). In contrast, pristine GQDs and *L*-His-GQDs caused only negligible decreases.

To determine whether membrane-order perturbation required a net nanochiral bias rather than histidine functionalization or surface charge alone, we introduced *DL*-His-GQDs prepared from racemic histidine. *DL*-His-GQDs exhibited a ζ-potential comparable to those of the enantiomeric His-GQDs but negligible CD signals, confirming the absence of a net ensemble chiroptical bias (Fig. S23). Flow-cytometric histograms showed that *D*-His-GQD treatment lowered the median Lo-associated NR12S fluorescence by approximately 32-fold relative to untreated WT-Spike cells, closely matching the shift produced by MβCD, whereas pristine GQDs, *L*-His-GQDs, and racemic *DL*-His-GQDs had little effect (Fig. 4e). In addition, *D*-His-GQDs tended to accumulate at cell–cell junctions in WT-Spike-expressing cells, a localization pattern that was largely abolished by MβCD pretreatment and restored by cholesterol repletion using cholesterol-loaded MβCD (MβCD–Chol) (Fig. 4f and S24). This spatial enrichment is consistent with enhanced interfacial interactions at protein-dense junctional membranes enriched in cholesterol- and sphingolipid-rich (raft-like) ordered nanodomains [58, 59]. Membrane damage associated with this interaction was assessed using LDH release and cell viability assays, which showed that pristine GQDs, *L*-His-GQDs, and *DL*-His-GQDs induced minimal LDH release and no appreciable loss of viability across the tested doses in WT-Spike- and Mut-Spike-expressing cells (Fig. 4g and h). By contrast, *D*-His-GQDs increased LDH release to 78.9% in WT-Spike cells and reduced viability in a concentration-dependent manner to 31.8% at 20 μg mL^-1^. Overall, this cell-based Spike display system demonstrates that WT Spike organizes ordered membrane nanodomains that serve as stereochemically addressable targets for nanochirality-dependent disruption by *D*-His-GQDs (Fig. 4i).

### 2.4. Biophysical basis of enantioselective GQD-membrane interactions

To elucidate how Spike S-acylation remodels the local viral membrane environment in ways that may enable recognition by nanoscale chirality, we performed all-atom molecular dynamics (MD) simulations of the Spike transmembrane domain (TMD) and cytosolic tail in a bilayer containing 1,2-dipalmitoyl-sn-glycero-3-phosphocholine (DPPC), 1,2-dilinoleoyl-sn-glycero-3-phosphocholine (DLiPC), and cholesterol [23]. Inspection of the atomistic snapshots revealed that the immediate annulus surrounding the S-acylated Spike became visibly depleted of the unsaturated lipid DLiPC, whereas DLiPC remained abundant around the non-acylated Spike (Fig. 5a). Quantification of the DLiPC mole fraction within the TMD inner region (≤2 nm) showed that, although both systems started from the same initial composition (37.5% DLiPC), the S-acylated trajectories converged to a much lower steady-state level (24.5%), whereas the non-acylated system remained close to the initial value (36.8%) (Fig. 5b). These observations indicate that palmitoylation promotes a more ordered local packing environment around Spike, in contrast to the DLiPC-rich, liquid-disordered (Ld)-like environment retained in the non-acylated system. Local organization within this acylation-associated packing field was assessed by quantifying the tilt angle, θ, between the principal lipid-tail axis and the local bilayer normal, n , in the inner and outer regions. Because smaller θ corresponds to a tail that is more aligned with n (i.e., more upright), the inner region exhibited a clear shift toward higher θ values, indicating more tilted tails in the Spike-proximal nanodomain relative to the outer membrane (Fig. S25). We next quantified the in-plane azimuthal angle, φ, of lipid-tail projections using 15° bins referenced to a fixed in-plane axis. The inner region showed stronger sectoral enrichment, indicating enhanced azimuthal anisotropy and a Spike-imprinted directional bias in local lipid packing (Fig. 5c and d). A mirror-contrast metric, Δ*K*_chiral_, was then used to quantify local chiral bias (Fig. 5e). In the S-acylated Spike system, Δ*K*_chiral_ was positive in the inner region (0.50) but remained near zero in the outer region (0.01), revealing a localized chiral bias in Spike-proximal lipid organization (Fig. 5f). Umbrella-sampling calculations further resolved the free-energy landscape associated with this stereoselective membrane engagement (Fig. 5g). At the respective PMF minima, local bilayer mapping placed *D*-His-GQDs approximately 0.85 nm beneath the local phosphate plane, compared with 0.51 nm for *L*-His-GQDs, indicating a deeper preferred insertion state for the *D* enantiomer (Fig. S26). Moreover, the potential of mean force (PMF) of *D*-His-GQDs increased substantially more steeply upon outward displacement from its minimum, indicating stronger stabilization of this deeper membrane-inserted state along the sampled reaction coordinate. These energetic and structural differences suggest that the Spike-proximal nanodomain provides a local basis for preferential *D*-His-GQD insertion and membrane engagement. The membrane-disruptive consequences of stereoselective insertion were then evaluated using vesicles with three lipid compositions spanning increasing raft-like character: PC, PC/cholesterol at 80:20, and PC/cholesterol/sphingomyelin at 40:40:20. *D*-His-GQDs reduced membrane order and induced time-dependent calcein leakage, with the strongest effect in the ternary raft-mimetic membrane (Fig. 5h and S27). TEM imaging similarly revealed greater *D*-His-GQD association with vesicle surfaces and more pronounced structural disruption in the more ordered membranes (Fig. 5i). Isothermal titration calorimetry (ITC) showed stronger apparent binding of *D*-His-GQDs to model membranes containing 40% cholesterol than of pristine GQDs, *DL*-His-GQDs, or *L*-His-GQDs, with an apparent association constant of (1.83 ± 0.31) × 10^6^ M^-1^ (Fig. 5j). This thermodynamic contrast independently reinforces the preferential association of *D*-His-GQDs with ordered membrane environments. Across these assays, structural chirality-encoded *D*-His-GQDs preferentially perturbed raft-like ordered membranes, in agreement with the Spike-associated nanodomain model (Fig. 5k).

**Fig. 5.**
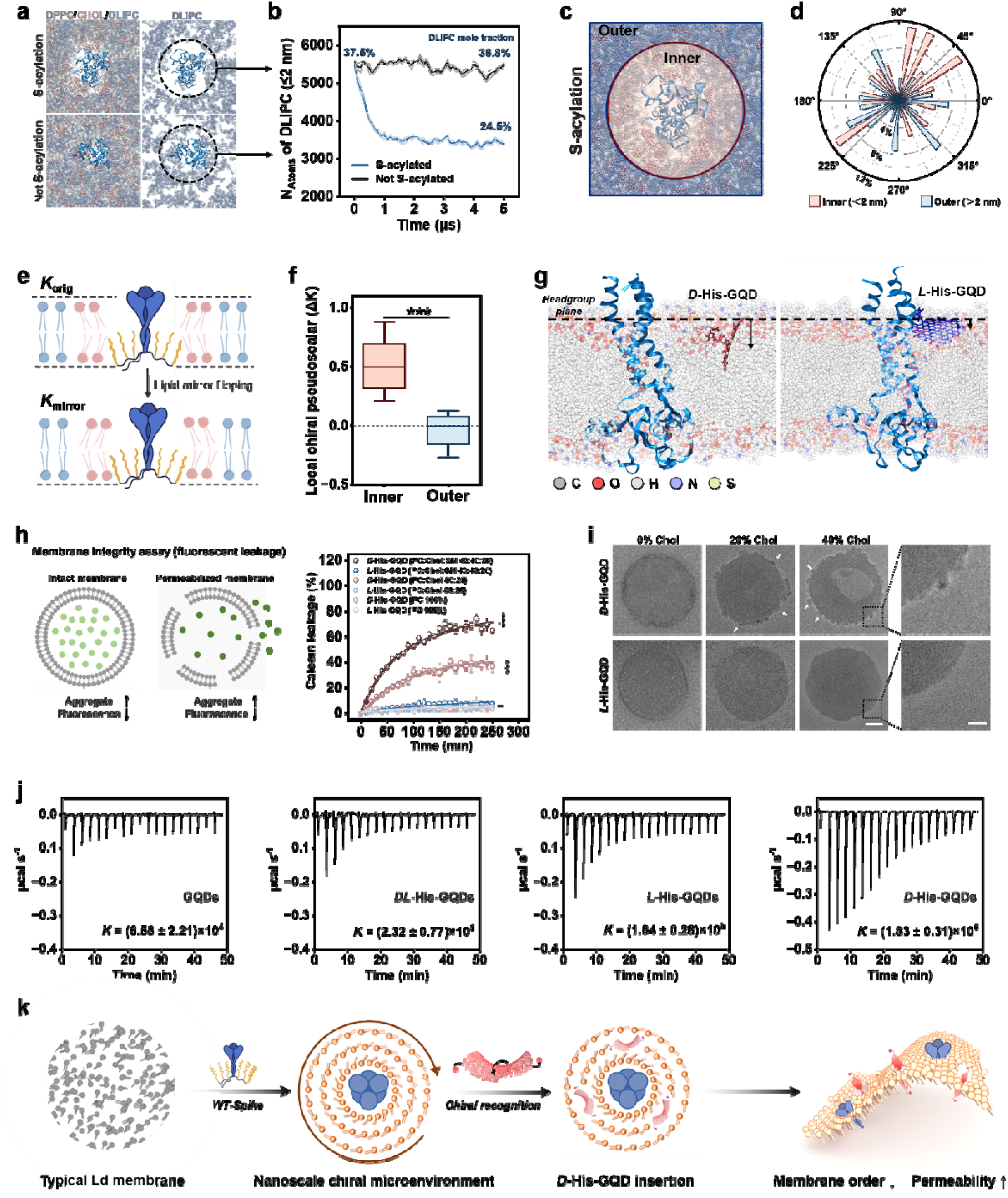
Molecular basis of stereoselective His-GQD insertion into ordered membranes. (a) Representative all-atom molecular dynamics (MD) snapshots of S-acylated and non-acylated Spike in a bilayer containing 1,2-dipalmitoyl-sn-glycero-3-phosphocholine (DPPC), 1,2-dilinoleoyl-sn-glycero-3-phosphocholine (DLiPC), and cholesterol (CHOL). (b) Time evolution of the number of DLiPC atoms within 2 nm of the Spike TMD. Annotated percentages indicate local DLiPC mole fractions calculated independently from lipid molecule counts. (c) Definition of the inner and outer membrane regions. (d) Polar distributions of lipid-tail azimuthal angles in both regions. (e) Mirror-contrast analysis used to calculate the local chiral pseudoscalar. (f) Local chiral pseudoscalar, Δ*K*_chiral_, in the inner and outer regions of the S-acylated Spike system (*n* = 5). Box plots show the median, quartiles, and range. (g) Representative MD snapshots of *D*- and *L*-His-GQDs interacting with the Spike-containing membrane. (h) Time-dependent calcein leakage from phosphatidylcholine (PC), PC/cholesterol (80:20), and PC/cholesterol/sphingomyelin (40:40:20) vesicles after treatment with *D*- or *L*-His-GQDs (n = 5). (i) TEM images of the corresponding vesicles. Scale bars, 100 nm in the main panels and 20 nm in the enlarged views. (j) Isothermal titration calorimetry thermograms of GQD interactions with model membranes. (k) Proposed model linking Spike S-acylation-associated membrane organization to stereoselective GQD interactions. Data in (h) are mean ± SD (*n* = 5). Statistical comparisons were performed using two-tailed *t* tests. \*\*\**P* < 0.001.

### 2.5. *In vivo* tolerability and antiviral protection against HCoV-OC43

To assess the *in vivo* feasibility of this nanochiral antiviral strategy, we first evaluated the short-term tolerability of intranasally administered His-GQDs (25 mg kg□^1^) in three-week-old C57BL/6 mice. Repeated administration of *L*- or *D*-His-GQDs once daily for three consecutive days did not adversely affect body-weight gain over the 8-day observation period (Fig. S28). Histological examination on day 8 revealed no evident treatment-related abnormalities in the kidney, heart, lung, liver, spleen, or brain (Fig. S29). Early organ distribution was assessed by ex vivo fluorescence imaging, which detected *L*- or *D*-His-GQDs-associated fluorescence in the lung and brain at 1 and 3 h after administration, with fluorescence approaching background by 6 h (Fig. 6a). Combined with the absence of treatment-related histological abnormalities, this transient profile indicates favorable short-term tolerability. We then evaluated antiviral efficacy using prophylactic and early post-exposure treatment schedules (Fig. 6b). Three-week-old C57BL/6 mice were challenged intranasally with 10 TCID of HCoV-OC43 and received intranasal *L*- or *D*-His-GQDs at 25 mg kg□^1^ either 1 h before or 1 h after viral challenge. PBS and remdesivir served as the vehicle and benchmark antiviral controls, respectively. Prophylactic administration produced a clear separation among groups. By day 7, body weight had fallen to 79% and 81% of baseline in mice receiving PBS and *L*-His-GQDs, respectively, whereas the *D*-His-GQD group reached 106%, closely matching uninfected controls (Fig. 6c). Protection was also evident with early post-exposure dosing (Fig. 6d). Mice given PBS or *L*-His-GQDs lost approximately 19–20% of their baseline weight by day 7, whereas *D*-His-GQDs limited the maximum loss to 5% and supported recovery to 108% by day 12, providing greater weight preservation than remdesivir. These body-weight trajectories demonstrate that *D*-His-GQDs provide substantial protection in both prophylactic and early post-exposure settings, whereas the *L*-enantiomer remains largely ineffective. Histopathological examination revealed extensive inflammatory-cell infiltration, alveolar septal thickening, and disrupted pulmonary architecture in the viral control, PBS, and *L*-His-GQD groups. In the brain, these groups exhibited increased parenchymal cellularity and focal accumulations of densely stained nuclei, consistent with infection-associated neuroinflammatory injury. *D*-His-GQDs markedly attenuated pulmonary and cerebral lesions and preserved tissue architecture close to that of normal controls (Fig. 6e). Representative CD3 immunofluorescence images showed stronger CD3 signals in the lung and brain of the viral control, PBS, and *L*-His-GQD groups, whereas lower signals were observed in the *D*-His-GQD group (Fig. 6f).

**Fig. 6.**
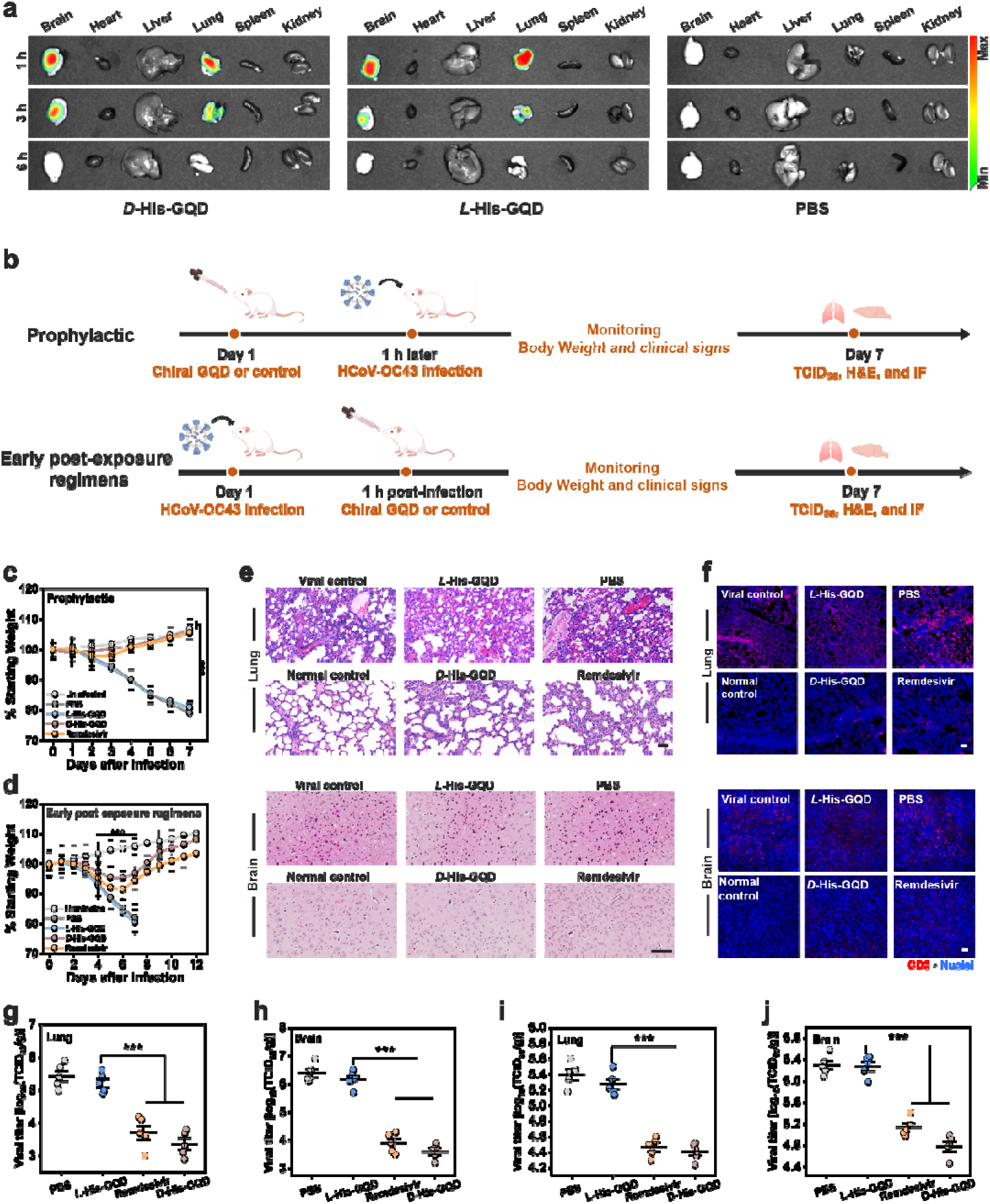
Distribution and antiviral efficacy of intranasal *D*-His-GQDs in HCoV-OC43-challenged mice. (a) Ex vivo fluorescence images of the brain, heart, liver, lung, spleen, and kidney at 1, 3, and 6 h after intranasal administration of *D*-His-GQDs, *L*-His-GQDs, or PBS. (b) Experimental timelines for prophylactic and early post-exposure administration. Changes in body weight relative to baseline in the prophylactic (c) and early post-exposure (d) studies. (*n* = 5). (e) Representative hematoxylin and eosin (H&E)-stained lung and brain sections. Scale bars, 50 μm. (f) Representative immunofluorescence images of lung and brain sections showing CD3□T cells in red and 4′,6-diamidino-2-phenylindole (DAPI)-stained nuclei in blue. Scale bars, 100 μm. Infectious viral titers in the lung (g) and brain (h) after prophylactic administration and in the lung (i) and brain (j) after early post-exposure administration. Titers are expressed as log_10_ 50% tissue culture infectious doses per gram of tissue (TCID_50_ g^-1^). (*n* = 5). Data are mean ± SD. Statistical comparisons were performed using two-tailed *t* tests. \*\*\**P* < 0.001.

Complementary immunofluorescence imaging showed lower viral Spike and CD45 leukocyte signals in the brain and lung following *D*-His-GQD treatment, while quantitative CD45 analysis confirmed lower leukocyte-associated fluorescence, consistent with reduced tissue inflammation (Fig. S30 and S31). TCID_50_ measurements quantitatively confirmed these protective effects. Prophylactic *D*-His-GQD administration lowered infectious viral titers by approximately 2.1 log in the lung and 2.7 log in the brain, whereas early post-exposure dosing achieved reductions of approximately 1.0 and 1.5 log in these tissues, respectively (Fig. 6g– j). *L*-His-GQDs showed little antiviral effect across either tissue or treatment schedule. Together, the body-weight, virological, histopathological, and immunological findings demonstrate that *D*-His-GQDs confer stereoselective protection when administered either before or shortly after HCoV-OC43 exposure.

## 3. Conclusion

This study identifies nanochirality as a material design parameter linking membrane interactions to antiviral selectivity. Histidine functionalization generates chiral GQD interfaces, with *D*-His-GQDs preferentially interacting with cholesterol-rich ordered membrane domains associated with S-acylated Spike. These interactions promote lipid disordering and membrane disruption, supporting direct extracellular virion inactivation. *D*-His-GQDs also reduce infection-associated syncytium formation. Compared with CPC and remdesivir, *D*-His-GQDs combine potent HCoV-OC43 inhibition with substantially higher cytotoxicity thresholds in uninfected human cell lines, demonstrating a favorable balance between antiviral activity and host-cell tolerance in the tested models. Stereoselective inhibition of HCMV in fibroblast and epithelial cells further supports the broader antiviral potential of this platform. Intranasal *D*-His-GQDs provide prophylactic and early post-exposure protection against HCoV-OC43 in mice, while repeated administration in uninfected animals shows favorable short-term tolerability. Together, these findings connect nanochiral interfacial architecture with preferential disruption of infection-associated membrane environments and support the development of antiviral biomaterials that combine membrane-directed activity with host-cell preservation.

## 4. Experimental section

### 4.1. Materials and instruments

Materials and reagents were purchased from commercial suppliers and used without further purification. All solvents were analytical pure unless otherwise noted. Distilled water was used throughout the experiments. Absorption spectra of liquid samples were determined on JASCO V-670 UV-Vis/NIR Spectrophotometer. The fluorescence spectra were recorded by an Infinite M1000 plate reader (Tecan). CD spectra were recorded on a CD spectroscopy (Jasco J-1700 Spectrometer). Transmission electron microscope (TEM) images were obtained using a FEI Talos F200S instrument. The zeta potentials were recorded using Malvern Zetasizer Nano ZS (Malvern Panalytical; Worcestershire, United Kingdom). Confocal fluorescence imaging was performed with Nikon A1R MP multiphoton microscopy. Atomic Force Microscope (AFM) images were recorded using Jupiter XR AFM. In vivo imaging was recorded using an AMI HT optical imaging system (Advanced Molecular Imager High Throughput, Spectral Instruments Imaging, Tucson, AZ, USA). Isothermal titration calorimetry (ITC) experiments were carried out on a MicroCal PEAQ-ITC calorimeter (Malvern Panalytical Ltd., Malvern, UK).

### 4.2. Synthesis and characterization of chiral GQDs

GQDs were synthesized following our previously established protocol [49]. Carbon nanofibers (0.4 g) were dispersed in 40 mL of a sulfuric acid and nitric acid mixture (3:1, v/v). The suspension was sonicated for 2 h and mechanically stirred for 6 h at room temperature, followed by heating at 120 °C for 10 h. After cooling to room temperature, the reaction mixture was diluted with ice-cold deionized water, and the pH was adjusted to 8.0 with sodium hydroxide. The resulting GQDs were purified by dialysis against deionized water for 3 days using a 1 kDa molecular-weight-cutoff membrane. The final GQD concentration was adjusted to 1 mg mL□^1^. Histidine-functionalized GQDs were prepared through EDC/Sulfo-NHS-mediated amidation of the edge carboxyl groups of GQDs. Briefly, 20 μL of EDC solution (100 mM) was added to 2.5 mL of GQD solution (100 μM), and the mixture was stirred for 10 min. Sulfo-NHS solution (40 μL, 100 mM) was then added, followed by sonication in an ice-water bath for 40 min. The activated GQDs were transferred to a 1 kDa molecular-weight-cutoff centrifugal filter and washed three times with deionized water to remove excess EDC and Sulfo-NHS. Subsequently, 40 μL of L-histidine or D-histidine solution (100 mM) was added to the activated GQDs, and the reaction mixture was stirred for 2 h to obtain *L*-His-GQDs or *D*-His-GQDs, respectively. *DL*-His-GQDs were prepared under identical conditions using 40 μL of a 100 mM racemic histidine solution containing equimolar amounts of *L*- and *D*-histidine. Unreacted histidine was removed by dialysis against deionized water using a 1 kDa molecular-weight-cutoff membrane.

### 4.3. Cell culture and virus propagation

Human rhabdomyosarcoma (ATCC CCL-136), SH-SY5Y neuroblastoma (ATCC CRL-2266), and HepG2 hepatocellular carcinoma (ATCC HB-8065) cells were obtained from ATCC and cultured in DMEM or EMEM supplemented with 10% fetal bovine serum (FBS) and 1% penicillin-streptomycin at 37 °C with 5% CO . HCoV-OC43 (ATCC VR-1558) was propagated in RD cells at 33 °C. Viral titers were determined by the TCID (50% Tissue Culture Infectious Dose) assay using the Reed-Muench method. Human lung fibroblasts MRC-5 (ATCC CCL-171) and human retinal pigment epithelial cells ARPE-19 (ATCC CRL-2302) were maintained in DMEM supplemented with 10% FBS, L-glutamine, and penicillin–streptomycin at 37 °C under 5% CO . BADrUL131-Y4 CMV-GFP, an AD169-derived HCMV strain with repaired UL131, was kindly provided by Thomas Shenk of Princeton University. [57] HCMV infections were performed at 85–90% cell confluence using an MOI of 0.5.

### 4.4. HCoV-OC43 antiviral assays

Dose-Response Assay: RD cells were infected with HCoV-OC43 (MOI = 0.5) in the presence of varying concentrations of pristine GQDs, *L*-His-GQDs, *D*-His-GQDs, remdesivir, or CPC. After 48 h, viral titers in the supernatant were quantified by TCID . The half-maximal effective concentration (EC_50_) was calculated using non-linear regression analysis. Time-of-Addition Assay: Cells were treated with *D*-His-GQDs, remdesivir, or CPC (10 μg mL^-1^) at different intervals: “Full-time” (-1 h to 48 h), “Entry” (-1 h to 0 h), and “Post-entry” (+2 h to 48 h). Viral loads were quantified at 48 h post-infection. Virucidal Kinetic Assay: HCoV-OC43 virions (10^5^ PFU) were incubated with compounds (10 μg mL^-1^) at 37 °C for indicated durations. The mixture was then diluted 1000-fold to prevent drug carryover and added to RD cells to determine residual infectivity.

### 4.5. HCMV antiviral assays and cytocompatibility

For time-of-addition assays, MRC-5 and ARPE-19 cells were seeded in 12-well plates at 2 × 10 cells per well and infected with CMV-GFP at an MOI of 0.5 under three treatment schedules using 10 μg mL^-1^ D- or *L*-His-GQDs. For virus preincubation, CMV-GFP and His-GQDs were incubated together for 1 h at 37 °C before inoculation. For co-treatment, virus and His-GQDs were added simultaneously. For post-inoculation treatment, cells were infected for 1 h, washed with PBS, and subsequently cultured in His-GQD-containing medium. Uninfected cells and untreated infected cells served as negative and infection controls, respectively. GFP-positive infection was assessed at 48 and 72 hpi. For concentration-response assays, MRC-5 and ARPE-19 cells were seeded in 96-well plates at 3 × 10 cells per well and pretreated for 2 h with 0.781–100 μg mL^-1^ *D*- or *L*-His-GQDs. CMV-GFP was then added directly to the His-GQD-containing medium at an MOI of 0.5, and cells were incubated for 48 h. Cells were fixed with 1% paraformaldehyde, stained with 5 μg mL^-1^ DAPI, and imaged in six fields per well using a 10× objective. Infection was quantified in ImageJ from GFP-positive and DAPI-positive cell counts and normalized to untreated infected controls. IC values were determined by linear interpolation between adjacent mean responses bracketing 50% relative infection on the log concentration scale. Cytocompatibility was evaluated using the Cell Meter Colorimetric Cell Cytotoxicity Assay Kit. MRC-5 and ARPE-19 cells were seeded at 3 × 10 cells per well and exposed to 0.1, 1, 10, or 100 μg mL^-1^ *D*- or *L*-His-GQDs for 24 h. Assay solution was then added at 20 μL per well, and absorbance at 570 and 605 nm was measured after 3, 4, and 5 h incubation. Untreated cells and cell-free medium served as viability and background controls, respectively.

### 4.6. Transmission electron microscopy

HCoV-OC43 virions were concentrated through a 20% sucrose cushion at 100,000 × g for 2.5 h at 4 °C. Purified virions were incubated with pristine GQDs, *L*-His-GQDs, *D*-His-GQDs, remdesivir, or CPC at 10 μg mL^-1^ for 1 h at 37 °C. Samples were adsorbed onto glow-discharged carbon-coated copper grids, stained with 2% uranyl acetate, and imaged at 80 kV. Untreated virions processed in parallel served as controls.

### 4.7. Plaque reduction assay

HCoV-OC43 virions were preincubated with vehicle, pristine GQDs, *L*-His-GQDs, *D*-His-GQDs, remdesivir, or CPC at 10 μg mL^-1^ for 1 h at 37 °C. The mixtures and equivalent input PFU were adsorbed onto confluent RD-cell monolayers in six-well plates for 1 h. Cells were overlaid with maintenance medium containing 1% low-melting-point agarose and incubated at 33 °C for 4-5 days. Monolayers were fixed with 4% paraformaldehyde and stained with 0.1% crystal violet. Plaques were counted, and plaque-forming units were normalized to the untreated-virus control.

### 4.8. Cell-surface display and confocal microscopy

Mammalian expression plasmids encoding full-length WT and mutant HCoV-OC43 Spike were generated in the pcDNA3.4 backbone by GenScript using a human-codon-optimized sequence based on NCBI RefSeq YP_009555241.1. In the cytoplasmic-tail eight-Cys-to-Ala mutant (8C-A), residues C1320, C1321, C1322, C1325, C1329, C1333, C1336, and C1337 were replaced with alanine. The corresponding sequence segments spanning residues 1315–1340 were LLFFICCCTGCGTSCFKKCGGCCDDY for WT Spike and LLFFIAAATGAGTSAFKKAGGAADDY for 8C-A Spike. Residue numbering follows YP_009555241.1. The 8C-A construct was designed to reduce Spike S-acylation. Co-localization: RD cells were transfected with WT or 8C-A Spike plasmids using Lipofectamine 2000. At 48 h post-transfection, lipid rafts were labeled with NR12S. Cells were fixed with 4% PFA (without permeabilization) and stained with anti-Spike antibody followed by Alexa Fluor 488-conjugated secondary antibody. Images were captured using a confocal laser scanning microscope (CLSM). Flow cytometry: At 48 h after transfection with WT Spike, RD cells were treated with pristine GQDs, *L*-His-GQDs, *D*-His-GQDs, or MβCD. Untreated WT-Spike cells served as controls. Cells were washed with ice-cold PBS and stained with NR12S at 4 °C in the dark. After washing, cells were immediately analyzed using flow cytometer. Debris and cell aggregates were excluded based on forward- and side-scatter profiles, followed by singlet gating. NR12S fluorescence was recorded in the Lo-associated short-wavelength channel using identical acquisition settings for all samples. Fluorescence distributions were analyzed using FlowJo software.

### 4.9. HCoV-OC43 infection-induced syncytium assay

RD cells were infected with HCoV-OC43 at an MOI of 0.5 for 24 h and used as effector cells. After removal of the viral inoculum and washing with PBS. Then, infected RD cells treated with 10 μg mL^-1^ pristine GQDs, *L*-His-GQDs, *D*-His-GQDs, remdesivir, or CPC. After 24 h of co-culture, cells were fixed with 4% paraformaldehyde for 15 min. Plasma membranes were stained with DiD, nuclei were counterstained with DAPI, and images were acquired using a confocal laser-scanning microscope under identical settings.

### 4.10. Liposome interaction assay

Liposomes with varying molar ratios of cholesterol were prepared by thin-film hydration followed by extrusion through polycarbonate membranes. To evaluate membrane integrity disruption, **c**alcein was encapsulated within the liposomes during hydration. Unencapsulated calcein was removed by dialysis against PBS at 4 °C until no fluorescence was detected in the dialysate. Liposomes were incubated with *D*-His-GQDs, and fluorescence recovery (indicating leakage) was monitored using a fluorescence spectrophotometer. Complete lysis (*F*_max_) was achieved by adding 0.1% Triton X-100. The leakage percentage was calculated as: Leakage (%) = (*F* - *F*_0_) / (*F*max - *F*_0_) × 100%. To assess changes in lipid order and raft integrity, liposomes were labeled with NR12S. Fluorescence emission spectra were recorded, and the GP value was calculated using the equation: GP = (*I*_ordered_ - *I*_disordered_) / (*I*_ordered_ + *I*_disordered_), where I_ordered_ and I_disordered_ represent the fluorescence intensities at 560 nm and 630 nm, characteristic of the ordered (L_o_) and disordered (L_d_) phases, respectively.

### 4.11. Isothermal titration calorimetry

ITC measurements were performed using a MicroCal PEAQ-ITC calorimeter. The sample cell contained model liposomes at 20 μM, and the injection syringe contained pristine GQDs, *DL*-His-GQDs, *L*-His-GQDs, or *D*-His-GQDs at 300 μM. GQD samples were sequentially titrated into the liposome suspension under constant stirring. Corresponding dilution heats were subtracted from the raw data. The resulting binding isotherms were fitted using a one-set-of-sites model to obtain the apparent association constant (K), binding enthalpy (ΔH), and stoichiometry (N).

### 4.12. DFT calculations

Geometry optimizations were performed using density functional theory (DFT) at the B3LYP/6-31G(d) level with Grimme’s D3 dispersion correction. Vertical excitation energies were calculated by time-dependent DFT (TD-DFT) at the same level. All electronic structure calculations were carried out using Gaussian 16. The frontier molecular orbitals (HOMO and LUMO), electrostatic potential (ESP) distributions, and reduced density gradient (RDG) isosurfaces were analyzed using Multiwfn v3.8. Electron–hole distributions associated with the low-lying excited states were also generated in Multiwfn. Molecular structures and orbital isosurfaces were visualized using GaussView and VMD as appropriate.

### 4.13. Molecular dynamics simulations

All-atom molecular dynamics simulations were carried out using GROMACS 2022.1. The Spike transmembrane domain (TMD) together with the membrane-proximal cytosolic tail was embedded in a DPPC/DLiPC/CHOL ternary bilayer, and fully S-acylated and non-acylated Spike systems were constructed in parallel. Proteins and lipids were described by the CHARMM36m force field with compatible lipid parameters, and water was modeled using TIP3P. The topologies of *D*-His-GQD and *L*-His-GQD were generated using CGenFF. All systems were solvated in explicit water, neutralized, and supplemented with 0.15 M NaCl. After energy minimization, equilibration and production runs were performed in the NPT ensemble at 310 K and 1 bar under periodic boundary conditions. Long-range electrostatics were treated using the particle mesh Ewald method, and all bonds involving hydrogen atoms were constrained using LINCS. For membrane analysis, lipids within 2 nm of the Spike TMD were defined as the inner region, and the remaining membrane was defined as the outer region. The local DLiPC mole fraction was monitored to evaluate acylation-dependent lipid redistribution. Lipid orientational organization was analyzed using the tilt angle (θ) relative to the local bilayer normal and the azimuthal angle (φ) of the in-plane lipid-tail projection. Local chiral bias was quantified by a mirror-contrast analysis, in which the original lipid configuration was compared with its mirror-reflected counterpart to obtain the pseudoscalar descriptor ΔK_chiral. For GQD insertion simulations, *D*-His-GQD or *L*-His-GQD was initially positioned in the aqueous phase above a pre-equilibrated Spike-containing membrane. The reaction coordinate was defined as the separation along the bilayer normal between the GQD center of mass and a local membrane reference constructed from phospholipid headgroup phosphorus atoms using a cylindrical pulling geometry. Steered molecular dynamics was first performed along this coordinate to generate a continuous membrane-approach and insertion pathway. Umbrella sampling windows were subsequently extracted along the reaction coordinate and restrained using a harmonic biasing potential. The resulting biased distributions were combined using the weighted histogram analysis method (WHAM) to obtain the one-dimensional relative potential of mean force (PMF), with the minimum of each PMF profile set to zero. To prevent lateral displacement of the GQD during the distal-control simulations, weak harmonic restraints were additionally applied in the membrane plane while leaving motion along the bilayer normal governed by the umbrella coordinate. GQD insertion depth was quantified relative to the local phosphate plane rather than the global bilayer midplane to account for local membrane deformation around the Spike-containing nanodomain. Positive insertion depth denotes penetration of the GQD center of mass below the local phosphate headgroup plane toward the membrane interior. Structural analyses were performed using GROMACS utilities and in-house scripts, and molecular graphics were generated with VMD 1.9.4.

### 4.14. Ethical statement

Ethical statement All the animal experiments were performed according to the protocols evaluated and approved by the Institutional Animal Care and Use Committee of the University of Notre Dame. (Approval Number: 26-02-9873)

### 4.15. In vivo studies

Three-week-old C57BL/6 mice were used throughout the study. For tolerability assessment, mice received PBS, *L*-His-GQDs, or *D*-His-GQDs intranasally at 25 mg kg^-1^ once daily for three consecutive days. Body weight was monitored through day 8, when the kidney, heart, lung, liver, spleen, and brain were collected for hematoxylin and eosin staining. For organ-distribution analysis, mice received a single intranasal dose of His-GQDs at 25 mg kg^-1^. The major organs were collected at 1, 3, and 6 h and analyzed ex vivo using an AMI HT imaging system. For antiviral studies, mice were challenged intranasally with 10 TCID of HCoV-OC43. *L*-His-GQDs or *D*-His-GQDs were administered intranasally at 25 mg kg^-1^ either 1 h before infection or 1 h after infection. Control groups received PBS or remdesivir. Body weight and clinical signs were monitored daily. Lung and brain tissues were collected on day 7 for infectious-virus titration, histopathological examination, and immunofluorescence analysis. An additional post-exposure cohort was monitored through day 12 for body-weight recovery. Infectious viral titers were determined in RD cells by TCID assay and expressed as log TCID_50_ g^-1^ tissue. Tissue sections were stained with hematoxylin and eosin or immunolabeled for HCoV-OC43 Spike, CD3, and CD45, with nuclei counterstained using DAPI.

### 4.16. Statistical analysis

Statistical analysezs were performed using OriginPro 2025. Data are presented as mean ± SD unless otherwise indicated. The value of n represents independent biological experiments, liposome preparations, molecular dynamics replicas, or animals, as specified in the figure legends. HCoV-OC43 concentration-response and cytotoxicity data were fitted by four-parameter logistic regression using log10-transformed concentrations. HCMV IC_50_ values were determined by linear interpolation on the log_10_ concentration scale. Group comparisons were made using an unpaired two-tailed Student’s *t*-test. All tests were two-sided, with *P* < 0.05 considered significant.

## CRediT authorship contribution statement

**Yichen Liu:** Conceptualization, Investigation, Formal analysis, Writing – original draft. **Mahnoosh Maleki:** Investigation, Formal analysis. **Pilar Pérez-Romero:** Supervision, Writing – review & editing. **Yichun Wang:** Conceptualization, Supervision, Funding acquisition, Writing – review & editing.

## Acknowledgments

This work was supported by the NIH MIRA (NIH 1R35GM15608-01) and the NSF Career Award (NSF CBET-2337387). This research was funded in part by the BELS Supplemental Professional Development Award (Bioengineering and Life Sciences Initiative, University of Notre Dame; Grant No. 374553-28016). The TEM and confocal microscopy images were acquired at the Notre Dame Integrated Imaging Facility. ITC data were acquired at the Biophysics Instrumentation Core at the University of Notre Dame. Computational work was performed using the high-performance computing clusters at the Center for Research Computing at the University of Notre Dame. Mice were bred and maintained under specific pathogen-free conditions at the Freimann Life Science Center at the University of Notre Dame. We appreciate the support provided by these facilities.

## Data availability

All data supporting the findings of this study are available in the article and its Supplementary Information.

