## Supplementary material for "Nanochiral graphene quantum dots preferentially target virus-organized membrane states for host-sparing antiviral activity": SI

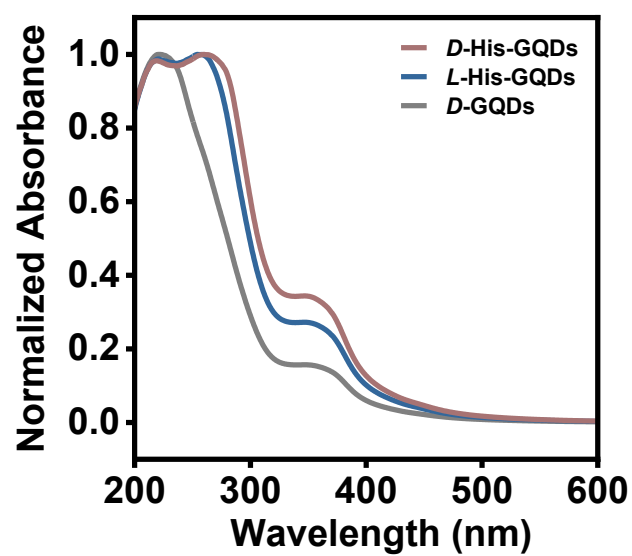

**Figure S1.** Normalized absorption spectra of GQDs, *L*-His-GQDs, and *D*-His-GQDs dispersed in PBS.

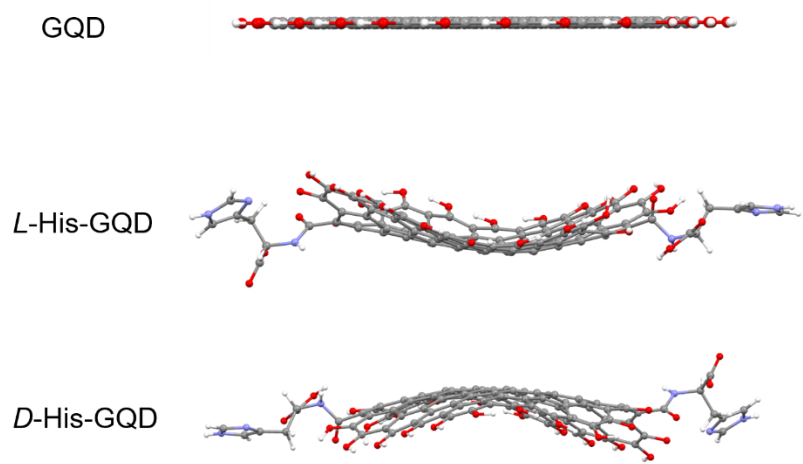

**Figure S2.** Optimized geometries of GQD, *L*-His-GQD, and *D*-His-GQD at the B3LYP/6-31G(d) level.

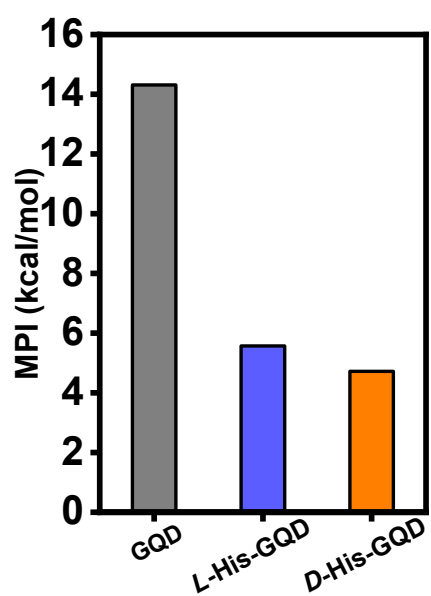

**Figure S3.** Molecular polarity indices (MPI) of GQD, *L*-His-GQD, and *D*-His-GQD.

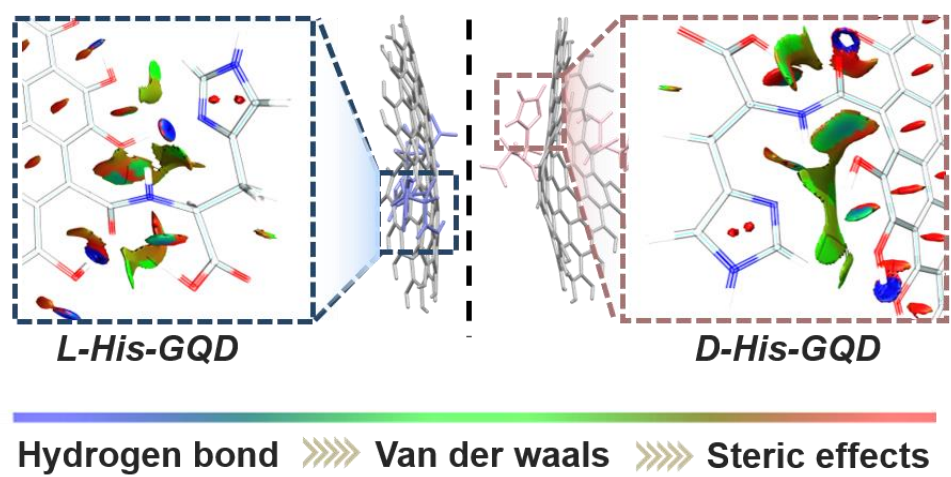

**Figure S4.** Reduced density gradient (RDG) isosurfaces of the *L*-His-GQDs and *D*-His-GQDs with RDG=0.5, colored based on the $\text{sign}(\lambda_2)\rho$.

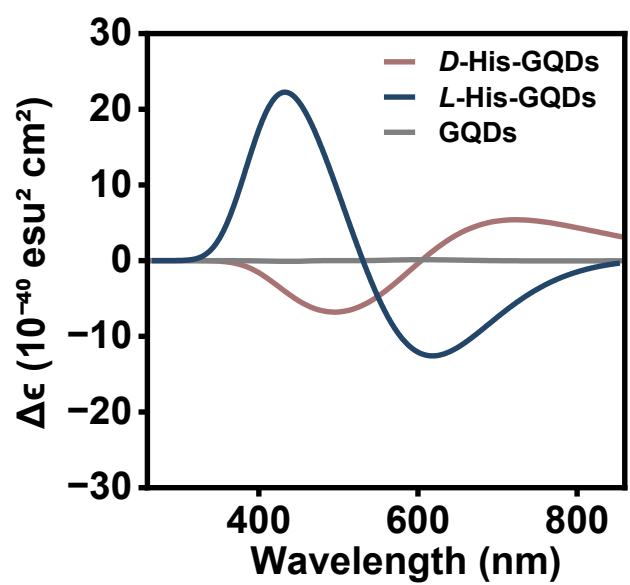

**Figure S5.** Simulated electronic circular dichroism (ECD) spectra (rotatory strength, $R \times 10^{-40}$ cgs) of GQD, *L*-His-GQD, and *D*-His-GQD calculated at the TD-B3LYP/6-31G(d) level; spectral lines obtained by Gaussian broadening with 0.25 eV HWHM.

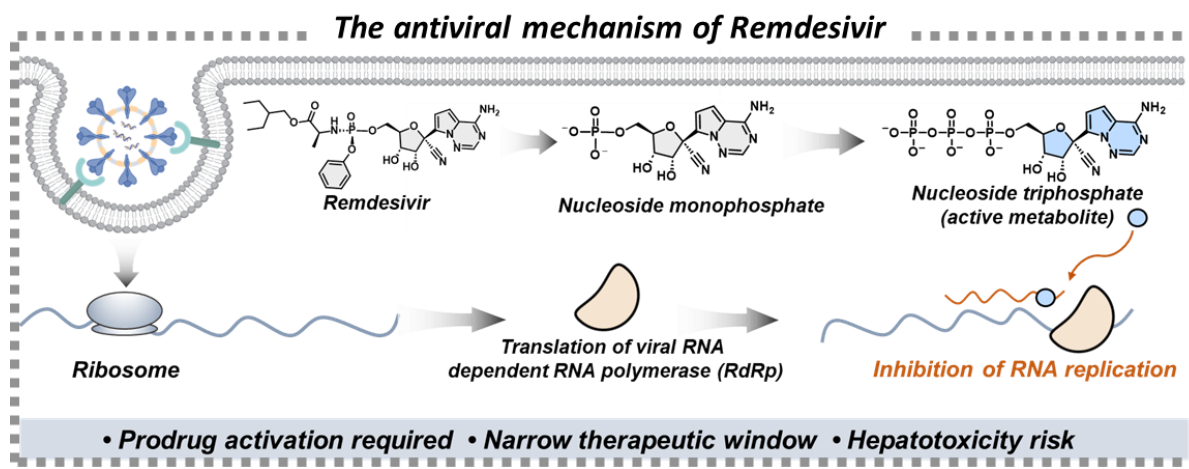

**Figure S6.** The antiviral mechanism of remdesivir.

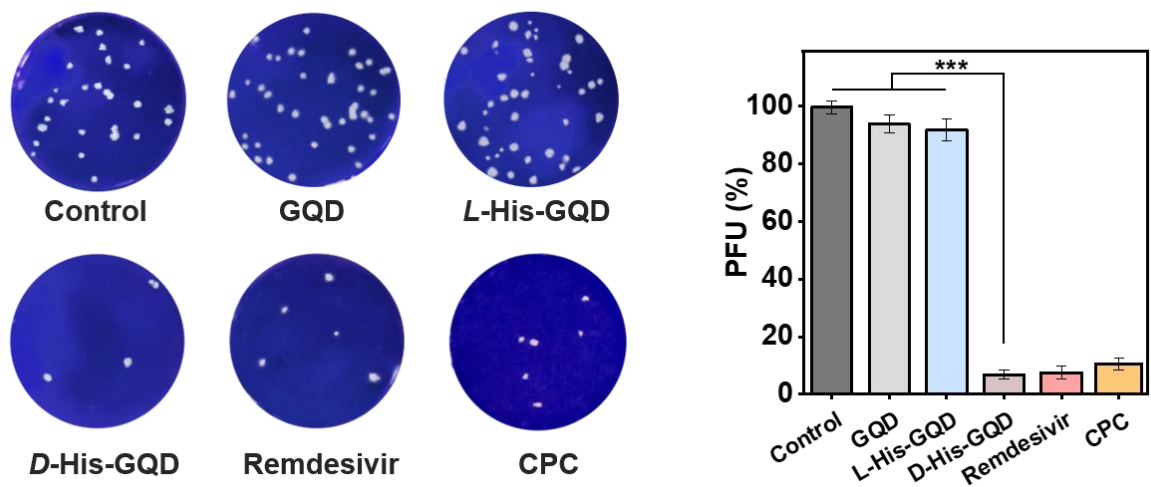

**Figure S7.** Infectious particles in the culture medium were detected by plaque formation. The histogram summarizes the plaque assay results. ($n = 5$). Data are expressed as the mean ± SD. Statistical differences were analyzed using two-tailed $t$ tests. \*\*\* $P < 0.001$.

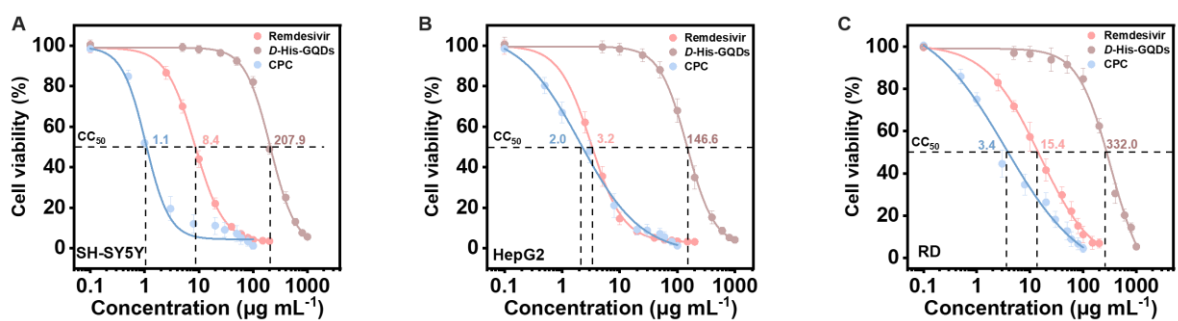

**Figure S8.** Cytocompatibility of *D*-His-GQDs, remdesivir and CPC. Dose-response viability curves in (A) SH-SY5Y neuronal cells, (B) HepG2 hepatocellular carcinoma cells, and (C) rhabdomyosarcoma (RD) cells treated with remdesivir, *D*-His-GQDs or CPC. Dashed lines mark CC<sub>50</sub> values. Concentrations are shown on a log scale.

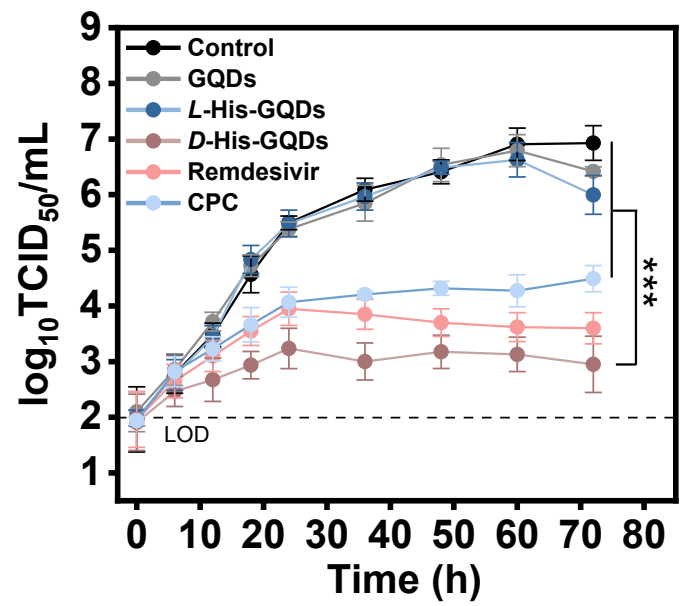

**Figure S9.** Time-dependent viral titers ($\log_{10}$ TCID<sub>50</sub>/mL) in the presence of Control, GQDs, *L*-His-GQDs, *D*-His-GQDs, remdesivir, or CPC. ($n=5$). Data are expressed as the mean ± SD. Statistical differences were analyzed using two-tailed $t$ tests. \*\*\*$P < 0.001$, \*$P < 0.05$.

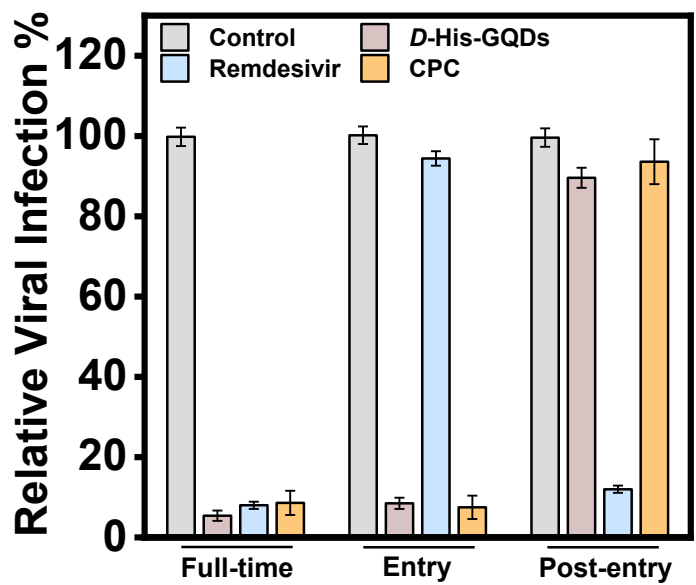

**Figure S10.** Stage-of-action profiles of *D*-His-GQDs, remdesivir, and CPC against HCoV-OC43. Relative infection was quantified after full-time, entry-only, or post-entry treatment. ($n=5$). Data are mean ± SD. Statistical differences were analyzed using two-tailed $t$ tests. \*\*\*$P < 0.001$

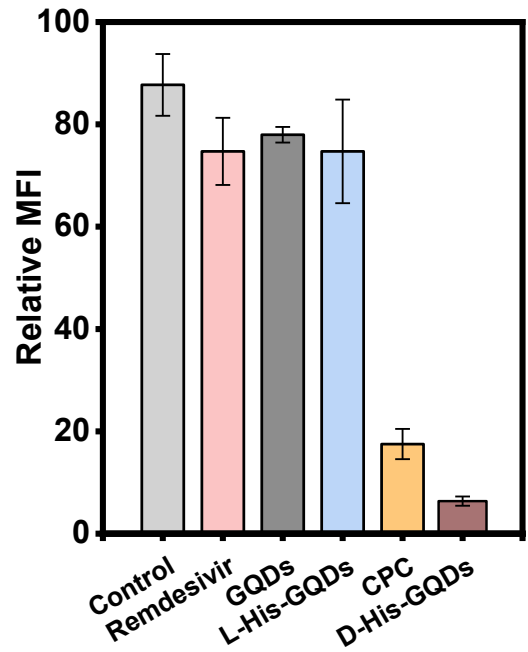

**Figure S11.** Mean fluorescence intensity (MFI) of the Spike signal was quantified in RD cells corresponding to the representative images in Fig. 3C. Data are mean ± SD from $n = 3$ independent experiments.

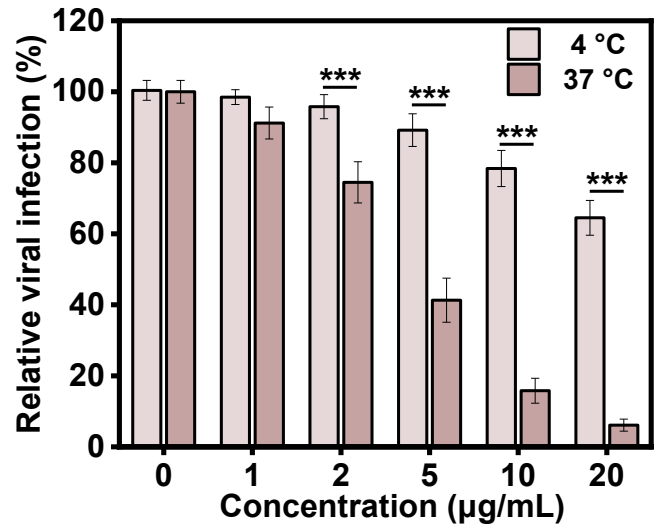

**Figure S12.** Temperature dependence of antiviral activity. Relative viral infection was measured after treatment at the indicated concentrations of *D*-His-GQDs at 4 °C or 37 °C. ($n = 5$) Data are mean ± SD. Statistical differences were analyzed using two-tailed $t$ tests. \*\*\*$P < 0.001$.

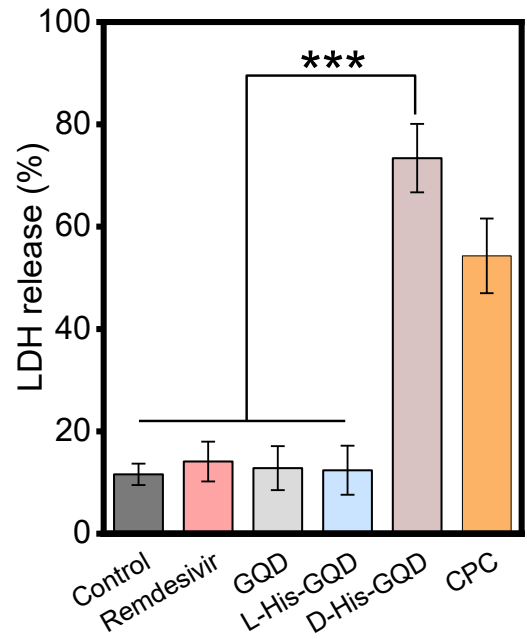

**Figure S13.** LDH release was measured for Control, remdesivir, pristine GQD, *L*-His-GQD, *D*-His-GQD, and CPC treatments. ($n = 5$) Data are mean ± SD. Statistical differences were analyzed using two-tailed $t$ tests. \*\*\*$P < 0.001$.

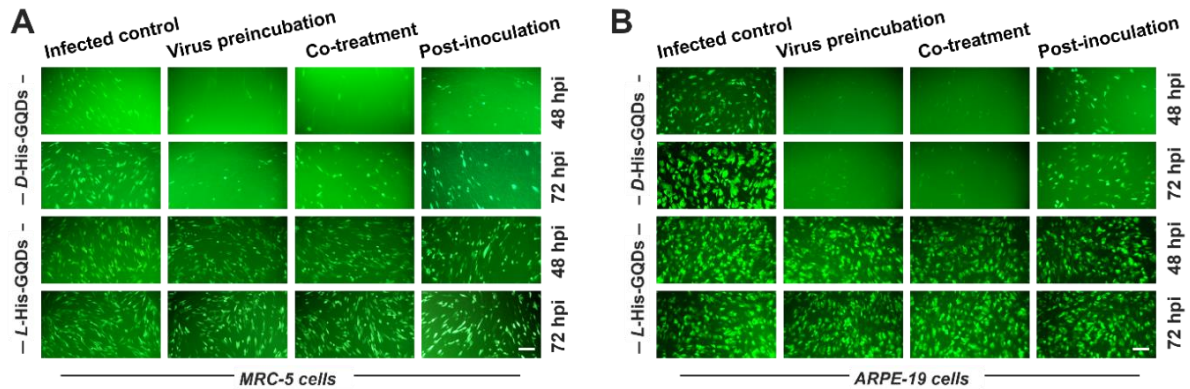

**Figure S14.** Enantioselective inhibition of human cytomegalovirus (HCMV) by *D*-His-GQDs. Representative green fluorescent protein (GFP) images of infected MRC-5 (**A**) and ARPE-19 (**B**) cells treated with $10 \mu\text{g mL}^{-1}$ *L*- or *D*-His-GQDs under the indicated schedules at 48 and 72 hours post-infection (hpi). Scale bars, $20 \mu\text{m}$.

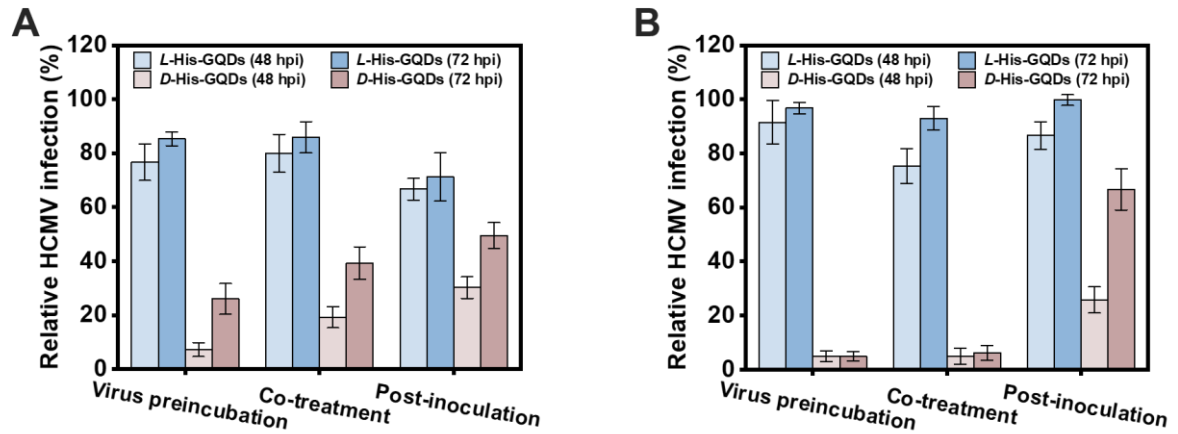

**Figure S15.** Time-of-addition analysis of HCMV infection in MRC-5 (A) and ARPE-19 (B) cells treated with $10 \mu\text{g mL}^{-1}$ D- or L-His-GQDs. Relative infection was measured at 48 and 72 hpi and normalized to untreated infected controls. Data are mean ± SD, $n = 4$.

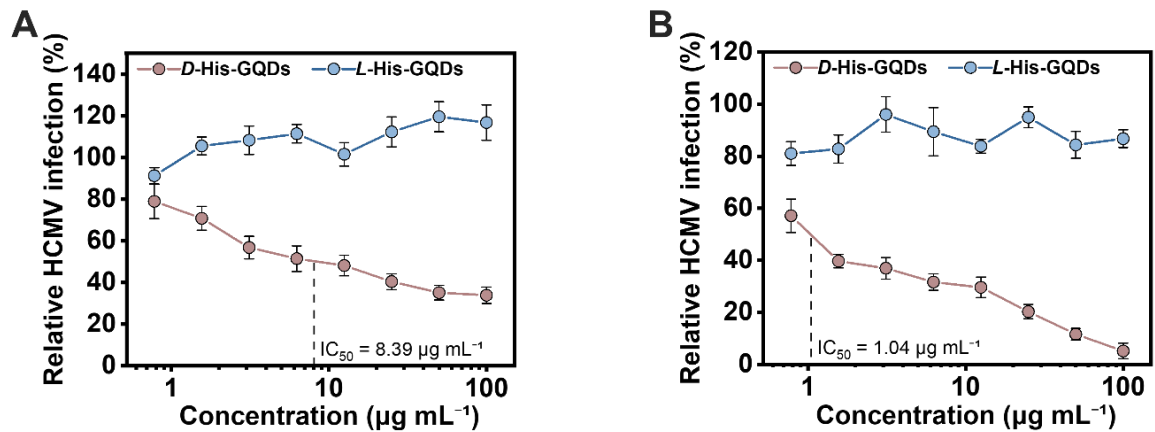

**Figure S16.** Concentration-dependent inhibition of HCMV infection in MRC-5 (A) and ARPE-19 (B) cells after 2 h of pretreatment followed by 48 h of infection in the continued presence of His-GQDs. Data are mean ± SD ($n = 4$).

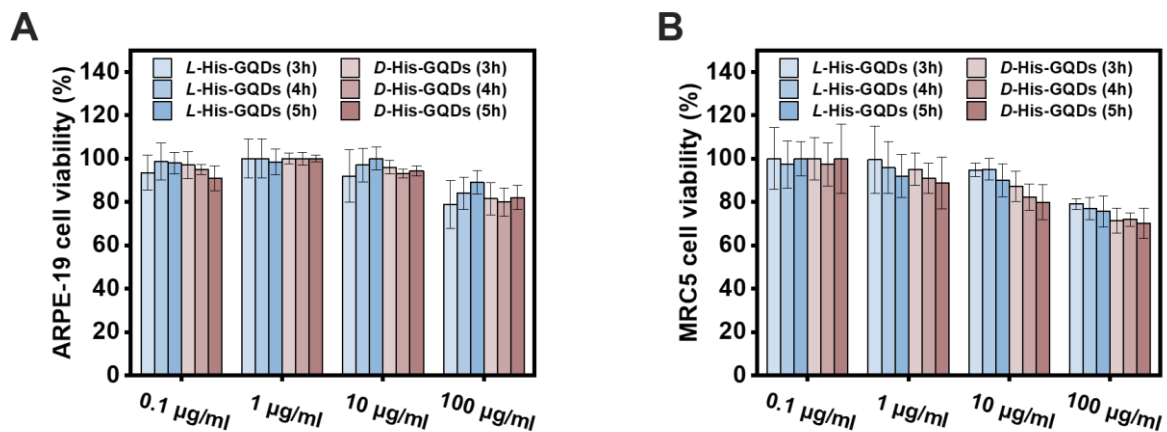

**Figure S17.** Effects of D- and L-His-GQDs on ARPE-19 (A) and MRC-5 (B) cell viability. Cells were treated with 0.1–100 $\mu\text{g mL}^{-1}$ His-GQDs for 24 h, and viability was measured after 3, 4, and 5 h incubation with the assay reagent. Data are mean ± SD, $n = 4$.

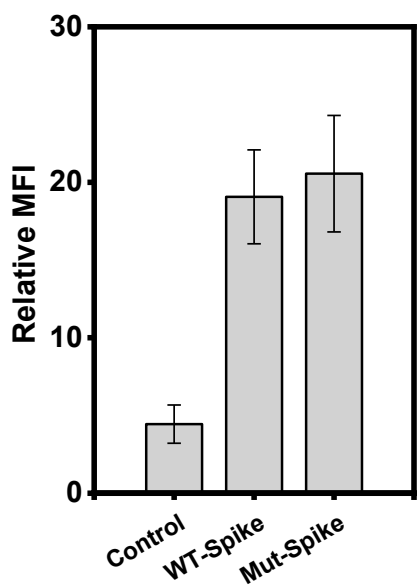

**Figure S18.** Cell-surface Spike MFI was quantified in non-permeabilized control, wild-type (WT)-Spike, and eight-Cys-to-Ala (8C-A)-Spike RD cells corresponding to Fig. 4B. Data are mean ± SD from $n = 3$ independent experiments.

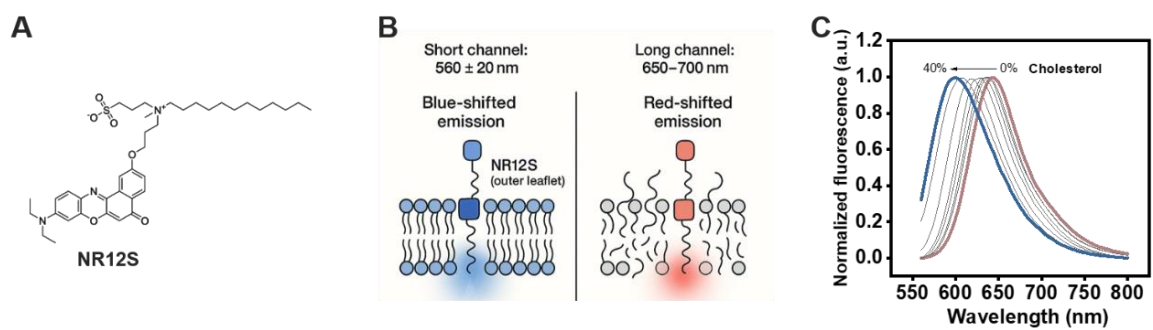

**Figure S19.** NR12S-based membrane order readout. **(A)** Chemical structure of the environment-sensitive probe NR12S. **(B)** Schematic of NR12S reporting lipid order through a blue-shifted short-wavelength channel and a red-shifted long-wavelength channel. **(C)** Representative normalized emission spectra showing a cholesterol-dependent blue shift with increasing membrane order.

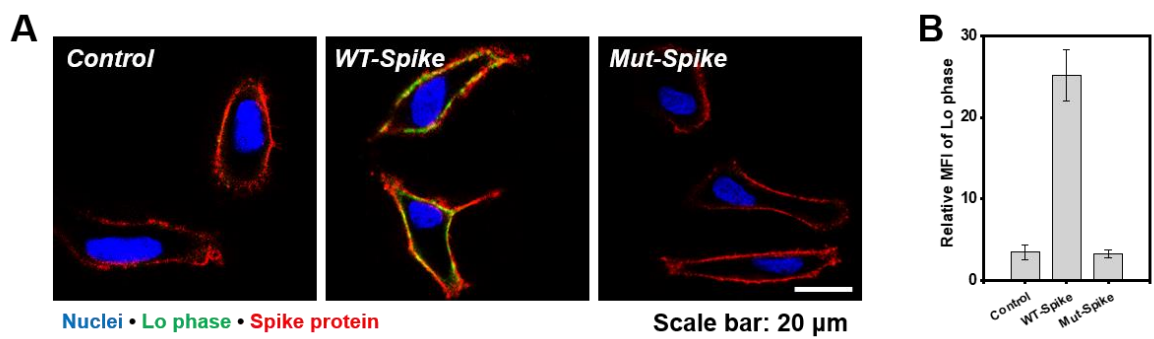

**Figure S20.** **(A)** Representative confocal images of control, WT-Spike, and 8C-A-Spike cells showing nuclei in blue, liquid-ordered (Lo)-associated NR12S fluorescence in green, and Spike in red. **(B)** Quantification of Lo-associated NR12S MFI. Scale bar, 20 μm. Data are mean ± SD from $n = 3$ independent experiments.

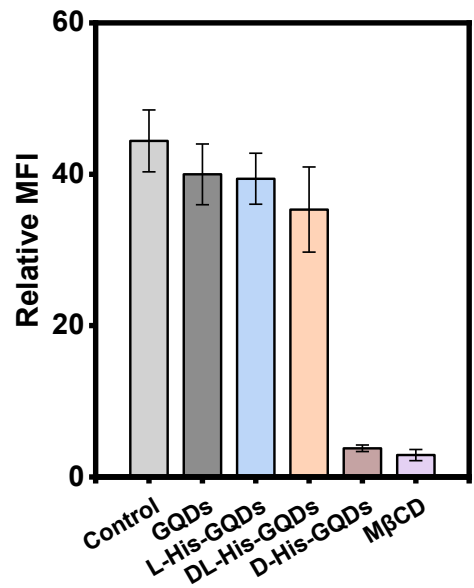

**Figure S21.** Lo-associated NR12S MFI was quantified at the 12 h endpoint of the imaging experiment shown in Fig. 4D after treatment with pristine GQDs, *L*-His-GQDs, *DL*-His-GQDs, *D*-His-GQDs, or methyl-β-cyclodextrin (MβCD). Data are mean ± SD from $n = 3$ independent experiments.

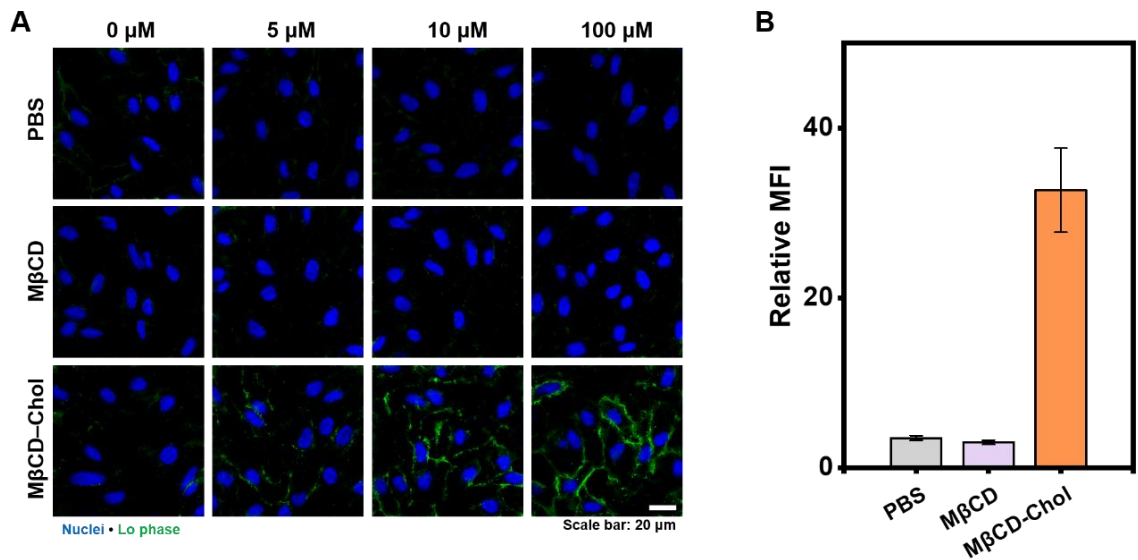

**Figure S22.** (A) Representative confocal images of *D*-His-GQD-pretreated WT-Spike cells subsequently exposed to PBS, MβCD, or cholesterol-loaded MβCD (MβCD-Chol) at the indicated concentrations. Nuclei and Lo-associated NR12S fluorescence are shown in blue and green, respectively. (B) Quantification of Lo-associated NR12S MFI at 100 μM. Scale bar, 20 μm. Data are mean ± SD from $n = 3$ independent experiments.

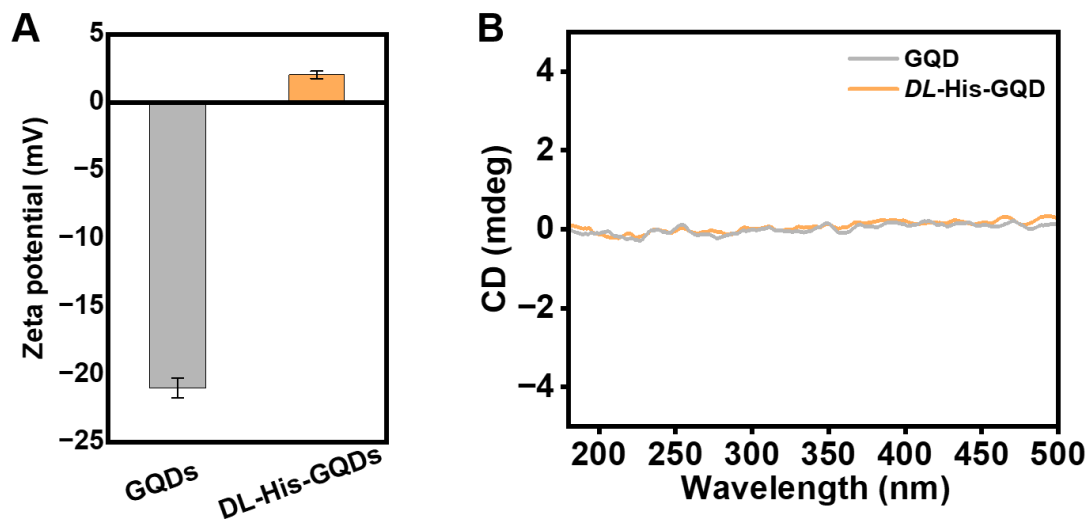

**Figure S23.** (A) ζ-potential of pristine GQDs and *DL*-His-GQD, showing charge modulation to a level comparable to chiral His-GQDs. Data are shown as mean ± SD ($n = 3$). (B) CD spectra of GQDs and *DL*-His-GQD.

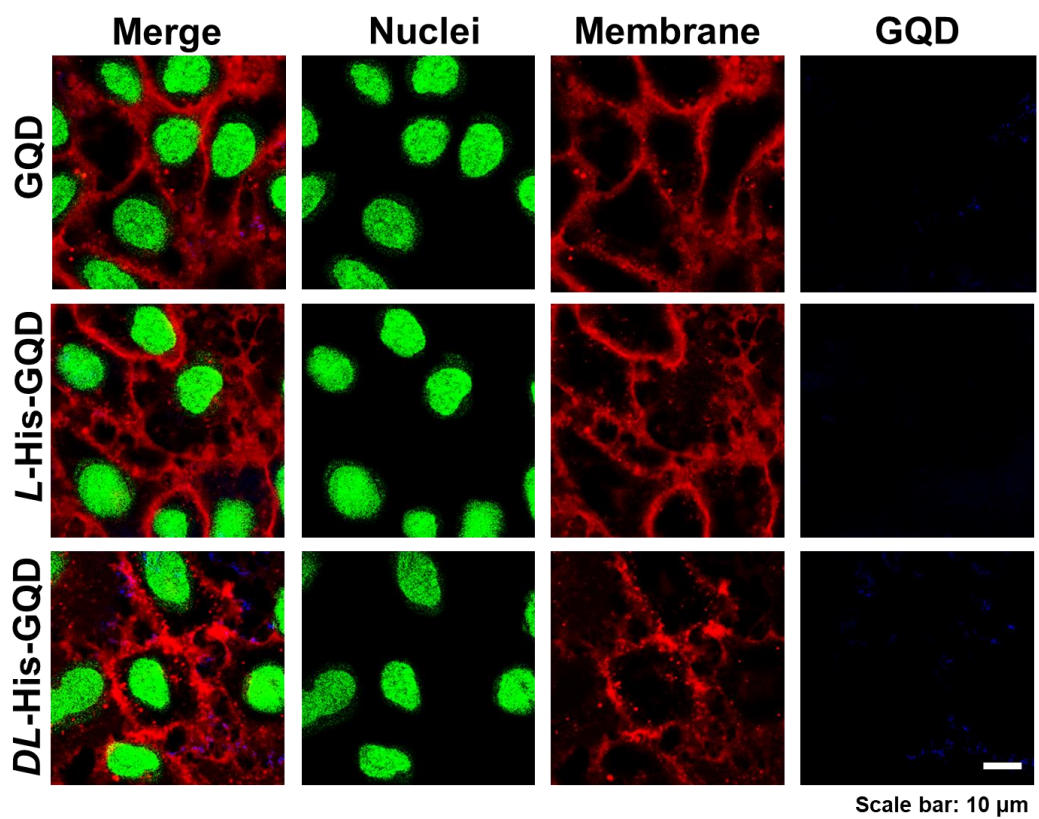

**Figure S24.** Confocal images of indicated GQD fluorescence in WT-Spike cells. Scale bar: 10 $\mu\text{m}$.

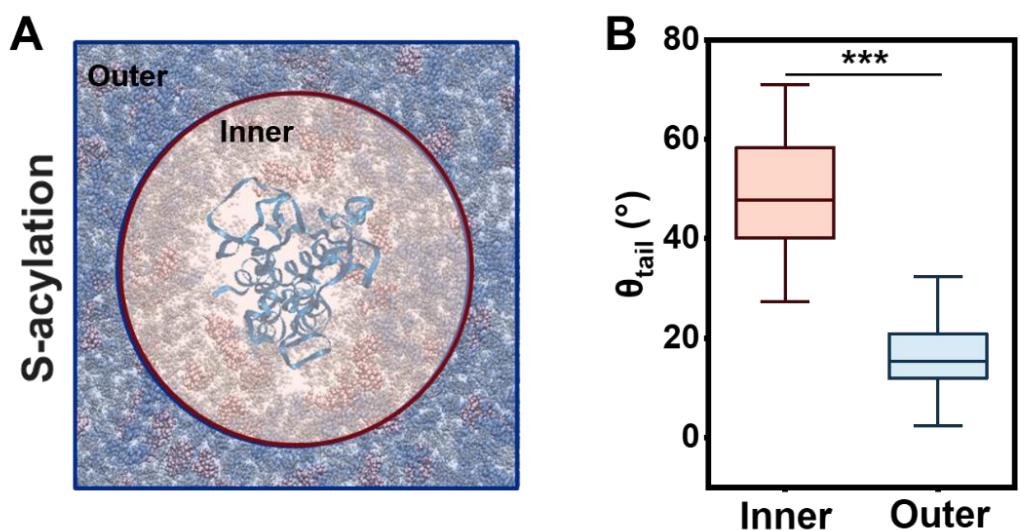

**Figure S25.** Spike-proximal nanodomain definition and lipid-tail tilt analysis. (A) Representative simulation snapshot illustrating the protein-centered inner region ($r < 2$ nm, pink shaded region) and the outer region ($r > 2$ nm, blue shaded region) used for local lipid analysis. (B) Box plots of lipid tail tilt angle $\theta_{tail}$ for inner and outer regions relative to the local bilayer normal. Box plots show the median and interquartile range with $1.5 \times \text{IQR}$ whiskers. ($n = 10$). Data are expressed as the mean ± SD. Statistical differences were analyzed using two-tailed $t$ tests. \*\*\*$P < 0.001$.

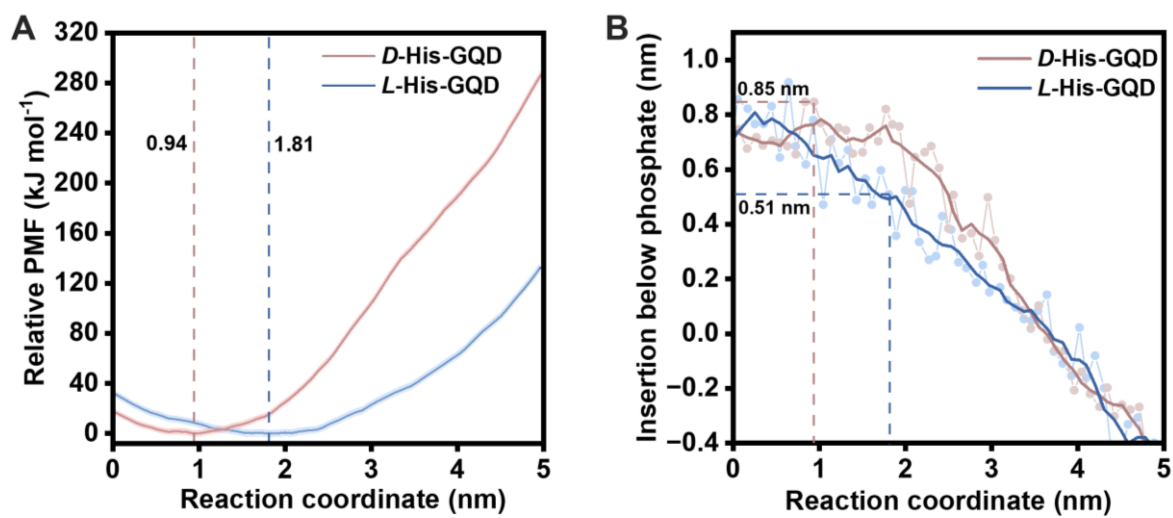

**Figure S26.** (A) Relative potential of mean force (PMF) profiles of *D*-His-GQD and *L*-His-GQD along the reaction coordinate; dashed lines indicate the respective PMF minima. (B) Insertion depth below the local phosphate plane, showing deeper preferred insertion of *D*-His-GQD (0.85 nm) than *L*-His-GQD (0.51 nm). Pale symbols denote individual windows, and solid lines indicate five-window moving averages.

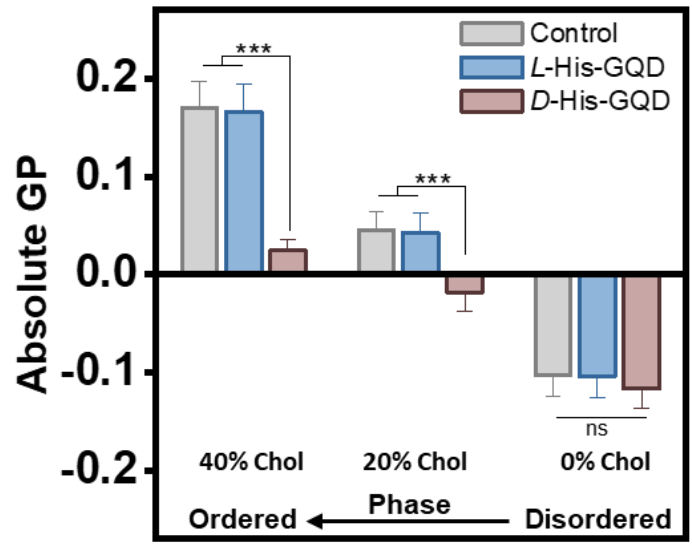

**Figure S27.** Lipid-composition-dependent perturbation of membrane order. Generalized polarization values were measured in PC, PC/cholesterol (80:20), and PC/cholesterol/sphingomyelin (40:40:20) bilayers after treatment with *L*- or *D*-His-GQDs. ($n = 5$). Data are mean ± SD. Statistical differences were analyzed using two-tailed $t$ tests. \*\*\*$P < 0.001$.

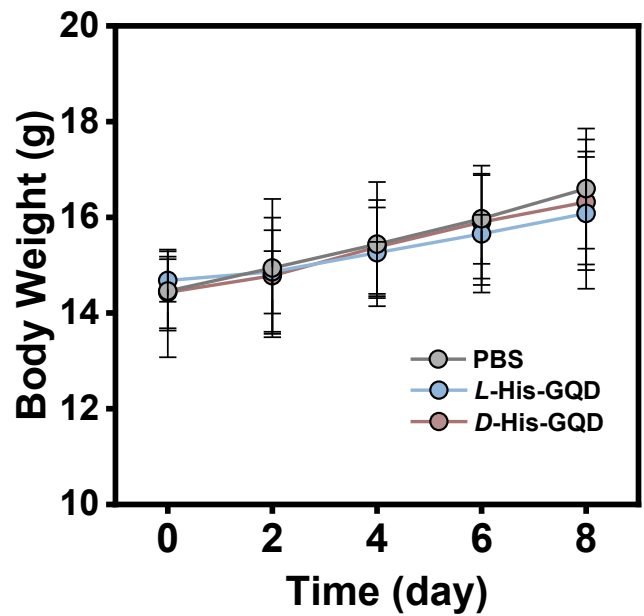

**Figure S28.** Body-weight trajectories of three-week-old C57BL/6 mice following intranasal administration of PBS, *L*-His-GQDs, or *D*-His-GQDs once daily for three consecutive days. ($n = 5$). Data are presented as mean ± SD.

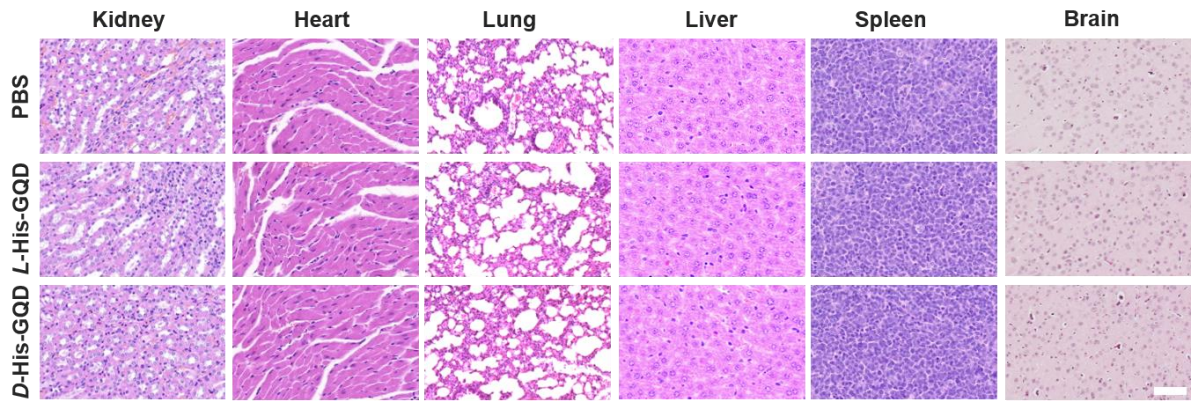

**Figure S29.** Three-week-old C57BL/6 mice received PBS, *L*-His-GQDs, or *D*-His-GQDs intranasally once daily for three consecutive days. Representative H&E-stained sections of the kidney, heart, lung, liver, spleen, and brain were collected on day 8. Scale bar, 50 μm.

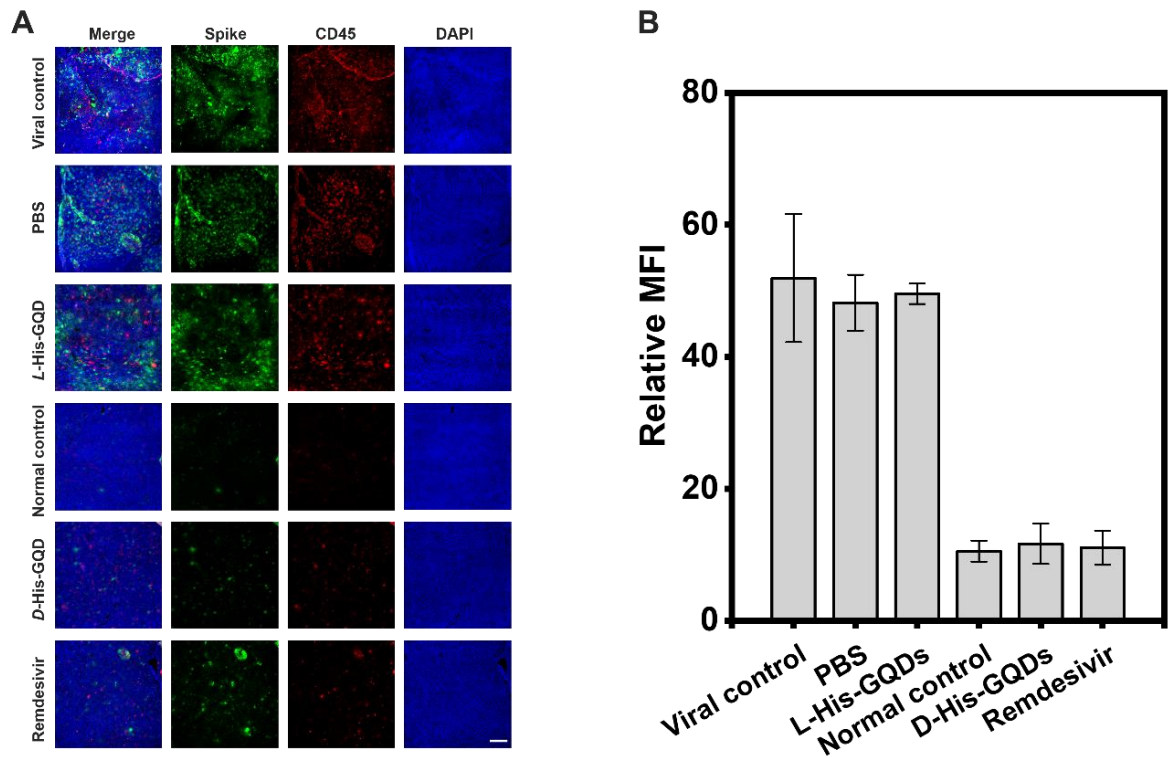

**Figure S30.** (A) Representative images of brain sections from the viral-control, PBS, *L*-His-GQD, normal-control, *D*-His-GQD, and remdesivir groups. HCoV-OC43 Spike, CD45<sup>+</sup> leukocytes, and DAPI-stained nuclei are shown in green, red, and blue, respectively. (B) Quantification of the relative mean fluorescence intensity (MFI) of the CD45 immunofluorescence signal. Scale bar, 100 μm. Data are mean ± SD from $n = 3$ mice per group, with multiple fields averaged for each animal.

**Figure S31.** (A) Representative images of lung sections from the viral-control, PBS, *L*-His-GQD, normal-control, *D*-His-GQD, and remdesivir groups. HCoV-OC43 Spike, CD45<sup>+</sup> leukocytes, and DAPI-stained nuclei are shown in green, red, and blue, respectively. (B) Quantification of the relative mean fluorescence intensity (MFI) of the CD45 immunofluorescence signal. Scale bar, 100 μm. Data are mean ± SD from $n = 3$ mice per group, with multiple fields averaged for each animal.
